# Haplotype-Resolved Long-Read Sequencing in Hundreds of Diverse Brains Identifies Structural Variant Impacts on Expression and Allele-Specific Methylation

**DOI:** 10.1101/2024.12.16.628723

**Authors:** Melissa Meredith, Kensuke Daida, Abraham Moller, Pilar Alvarez Jerez, Shloka Negi, Laksh Malik, Rylee M. Genner, Xinchang Zheng, Sophia B. Gibson, Mira Mastoras, Breeana Baker, Cedric Kouam, Kimberly Paquette, Paige Jarreau, Mary B. Makarious, Anni Moore, Samantha Hong, Dan Vitale, Syed Shah, Jean Monlong, Caroline B. Pantazis, Mobin Asri, Kishwar Shafin, Paolo Carnevali, Stefano Marenco, Pavan Auluck, Ajeet Mandal, Karen H. Miga, Arang Rhie, Xylena Reed, Jinhui Ding, Andrew Singleton, Danny E. Miller, Mark Chaisson, Winston Timp, J. Raphael Gibbs, Mark R. Cookson, Adam M. Phillippy, Mikhail Kolmogorov, Fritz J. Sedlazeck, Miten Jain, Mike Nalls, Benedict Paten, Cornelis Blauwendraat, Kimberley J. Billingsley

## Abstract

Structural variants (SVs) drive gene expression in the human brain and are causative of many neurological conditions. However, most existing genetic studies have been based on short-read sequencing methods, which capture fewer than half of the SVs present in any one individual. Long-read sequencing (LRS) enhances our ability to detect disease-associated and functionally relevant structural variants; however, its application in large-scale genomic studies has been limited by challenges in sample preparation and high costs. Here, we leverage a new scalable wet-lab protocol and computational pipeline for whole-genome Oxford Nanopore Technologies sequencing and apply it to neurologically normal control samples from the North American Brain Expression Consortium (NABEC) (European ancestry) and Human Brain Collection Core (HBCC) (African or African admixed ancestry) cohorts. Through this work, we present a publicly available long-read resource from 351 human brain samples (median N50: 27 Kbp and at an average depth of ∼40x genome coverage). We discover approximately 234,905 SVs and produce locally phased assemblies that cover 95% of all protein-coding genes in GRCh38. To resolve cis-regulatory effects, we develop ASM-LR, a method for allele-specific methylation analysis from long-read data, revealing both strong and subtle regulatory effects, including numerous novel methylation QTLs masked in unphased models. Our results highlight the power of haplotype-resolved methylation to uncover regulatory mechanisms and establish a foundational resource for exploring how genetic variation shapes gene expression and epigenetic architecture across diverse ancestries.

## Main

Structural variants (SVs), including insertions, deletions, duplications, inversions, and complex rearrangements, represent significant contributors to genetic variation within the human genome. These variations can influence gene expression, regulatory elements, and genomic stability, playing a critical role in shaping the genetic architecture of the human brain. Through these mechanisms, SVs impact individual brain structure^1^, cognitive abilities, and susceptibility to a range of neurological disorders ^2,3^. Notably, SVs are implicated in the pathogenesis of several neurodegenerative diseases. For instance, duplications of the *APP* gene have been linked to early-onset Alzheimer’s disease (AD)^4^, while duplications and triplications of the *SNCA* gene contribute to Parkinson’s disease (PD) and dementia with Lewy bodies (DLB) ^5^. Similarly, transposable element insertions in the gene *TAF1* are causative of X-linked dystonia Parkinsonism ^6–8^, and *PRKN* (Parkin) gene rearrangements are a known cause of early onset PD^9–12^. These examples underscore the critical role of SVs in the etiology of neurodegenerative disease and highlight the need for comprehensive characterization of these variants to fully understand brain disorders.

Despite their importance, conventional short-read sequencing (SRS) methods, while useful for many applications, are limited in their ability to accurately detect and genotype SVs^13^. The short read lengths, typically only a few hundred base pairs, make it difficult to span larger SVs, especially those in the kilobase range. Additionally, SRS struggles to resolve variants within repetitive genomic regions, which hinders both the resolution of SVs and the accurate phasing of variants, leading SRS-based approaches to miss more than half of the SVs present in any one individual. Long-read sequencing (LRS) technologies address these limitations by providing longer reads that can more effectively span large SVs, resolve repetitive regions, and phase variants across much larger genomic distances. Recent studies, including our own, have shown that LRS significantly improves SV detection and variant phasing, even in challenging genomic regions^14,15^.

Beyond these technical limitations, the historical focus on European populations in genomic research has also significantly restricted our understanding of genetic variation in other ancestries. This bias has hindered the discovery of population-specific variants, which are essential for understanding traits such as disease susceptibility, drug metabolism, and gene regulation across different populations. For example, the power of diverse datasets was demonstrated when a population-specific *GBA1* variant associated with PD was identified in African or African admixed ancestry populations^16^. As research continues to reveal substantial differences in genetic risk factors across ancestries, the development of inclusive and comprehensive genomic resources becomes crucial for improving the accuracy of disease mapping^17^ and ensuring more equitable, globally relevant healthcare solutions.

In this study, we generate high-quality long-read whole-genome sequencing data from the postmortem brains of 351 neurologically normal individuals of European and African or African admixed ancestry. This effort results in the discovery of 234,905 SVs and the generation of locally phased assemblies covering 95% of all protein-coding genes in GRCh38 (98% of single-copy genes). By integrating these data with matched transcriptomic and methylation profiles we perform quantitative trait locus (QTL) analyses, and identify numerous SVs that influence gene expression and DNA methylation in the human frontal cortex. To enable haplotype-resolved analysis of epigenetic regulation, we developed ASM-LR, a novel tool for detection of allele-specific DNA methylation from long-read sequencing data (https://github.com/molleraj/CARDlongread_allele_specific_QTL). The full dataset is available through the AnVIL platform with dbGaP restricted access (NABEC: https://explore.anvilproject.org/datasets/a00842c8-12f8-4f92-a7b3-01bb73ffd4e4 HBCC: https://explore.anvilproject.org/datasets/a4e936d1-d81a-475d-95be-b5cd41de921d) and the Alzheimer’s Disease Workbench (ADWB, https://www.alzheimersdata.org/ad-workbench). Together, these resources provide a foundational reference for understanding the regulatory impact of genetic variation in the brain and represent the first large-scale, ancestrally diverse control dataset with matched long- and short-read genomic and transcriptomic data.

## Results

### Genetic ancestry and sequencing overview of brain samples across cohorts

We conducted ONT whole-genome sequencing on 359 human brain samples from the NABEC (n=205) and HBCC (n=154) cohorts using the PromethION platform (Figure 1a). For the NABEC samples, sequenced on R9.4.1 flow cells using the LSK-110 library kit (referred to as R9), we obtained an average genome coverage of 46x and read N50 of 26.2 Kb. For the HBCC samples, sequenced on R10.4.1 flow cells and LSK-114 library kits (referred to as R10), we obtained an average genome coverage of 39x and read N50 of 24.6 Kb (Figure 1b, Supplementary Table 1). To determine genetic ancestry, NABEC (Illumina) and HBCC (R10 ONT) SNVs were evaluated using data from the 1000 Genomes project, the Human Genome Diversity Project, and an Ashkenazi Jewish panel with GenoTools^18,19^ (Supplementary Table 2). Comparative analysis with 1000 Genomes samples showed that, as expected, NABEC samples predominantly clustered with the European (EUR) group, while HBCC samples were more dispersed but predominantly clustered with the African ancestry cluster (Figure 1d). Eight individuals from the HBCC cohort, labeled as EUR, SAS, and AMR, were excluded from downstream analyses to reduce population structure and admixture effects in the QTL analysis. The final set of 351 samples was processed using the NAPU pipeline^14^, generating four key locally phased outputs for downstream analyses: de novo assemblies, read and assembly based SV calls, read-based small variant calls, and haplotype-resolved methylation calls.

**Figure 1:**
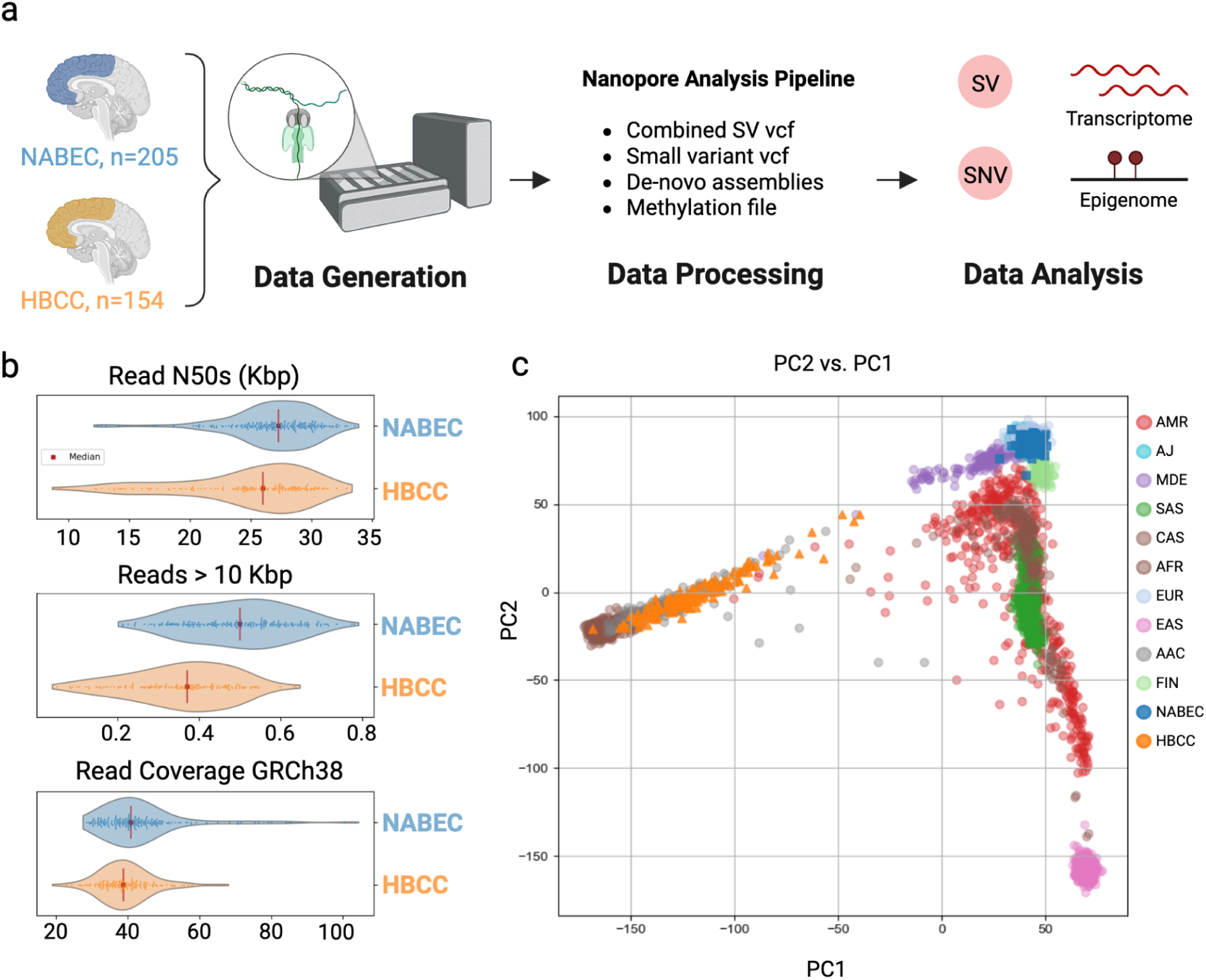
Overview of the study and samples. **a.** Graphical overview of the project workflow. **b.** Read N50s, proportion of reads longer than 10Kbp, and read coverage of GRCh38 for each cohort. Median values are denoted by the red vertical lines. **c**. Principal Component Analysis (PCA) of ancestry predictions using NABEC Illumina SNVs (blue squares) and HBCC ONT SNVs (orange triangles) overlaid on 1000 Genomes SNV ancestry clustering confirming that the NABEC cohort is predominantly of European ancestry while the HBCC cohort is of African or African admixed ancestry.

### Generating Genome Assemblies at scale

All samples were assembled with Shasta^20^ and phased into diploid assemblies with Hapdup^14^. Hapdup uses the ONT reads to phase Shasta assembly contigs and produce locally phased ‘dual’ and phased assemblies, in which contigs are the phase blocks. Shasta was run with an R9 and R10 ONT configuration for the NABEC and HBCC cohorts respectively and produced dual assemblies with a median NG50 of 21.4 megabases (Mb) and 15.7 Mb (measured against the 3.1 gigabase (Gb) length of the T2T-CHM13v2.0 reference). The phase block median NG50 is 1.25 Mb and 2.43 Mb for NABEC and HBCC, respectively. NG50s are lower for HBCC dual assemblies but, due to the heterozygosity of the African or African admixed HBCC samples, we observed longer phase blocks for HBCC assemblies compared to the European ancestry NABEC samples (Supplementary Figure 1a, 1b). Despite these differences, samples from both cohorts assembled on average 94% of GRCh38 with a median 2.88 Gb assembly size (Supplementary Figure 1c). We found that Shasta assembled 99% of the regions labeled as “difficult” in GRCh38 by the Genome In a Bottle Consortium (GIAB)^21^. To measure assembly quality we estimated the base level accuracy by alignment to the CHM13 reference and computed a median divergence from the reference of 0.26-0.28% (contig identities to be 0.9971 and 0.9973) for NABEC and HBCC, respectively (Supplementary Figure 1d).

Next, we assessed the quality and usefulness of the assemblies’ gene models. Across both cohorts 97% of single copy genes are assembled, calculated by an asmgene gene-level analysis. The median number of protein-coding genes assembled completely in both haplotypes across all samples is 19,181 (95.6%); mean 19,048, (95.0%) (Supplementary Figure 1e). Shasta, however, struggled to assemble highly similar multicopy genes; on average ∼55% of those present in CHM13 were recovered in these assemblies. We examined 168 genes identified as associated with PD or AD in previous GWAS studies ^22,23^ (Supplementary Table 3). On average, 95% of the GWAS genes (160 of 168) are assembled contiguously. The majority of the genes reported as not completely assembled are, upon further inspection, partially assembled in one or both of the assembly haplotypes. Notably, we assembled 97% (169 median) of the 174 immunoglobulin heavy locus (IGH) genes in Genbank, which have been associated with AD and PD disease risk. Assembling the IGH loci in these 351 primary tissue samples is an important contribution of our work, as many large scale long read sequencing projects have predominately used lymphoblast cell lines.

### Structural variant characterization across diverse cohorts

We identified SVs using two complementary approaches: Sniffles^15^, a read alignment-based approach, and Hapdiff^14^, an assembly-based method. Outputs from both methods were merged using Truvari to produce a harmonized union callset. In that merged callset the majority of SVs were detected by both Sniffles and Hapdiff, 73% of SVs with allele frequency > 0.01 (Figure 2a). Some SVs, however, were only identified by just one method. Hapdiff in particular called more small insertions likely reflecting its’ assembly-based approach (Figure 2c). Most SVs were rare, with 59% having a minor allele frequency (MAF) below 1% (Figure 2d).

**Figure 2:**
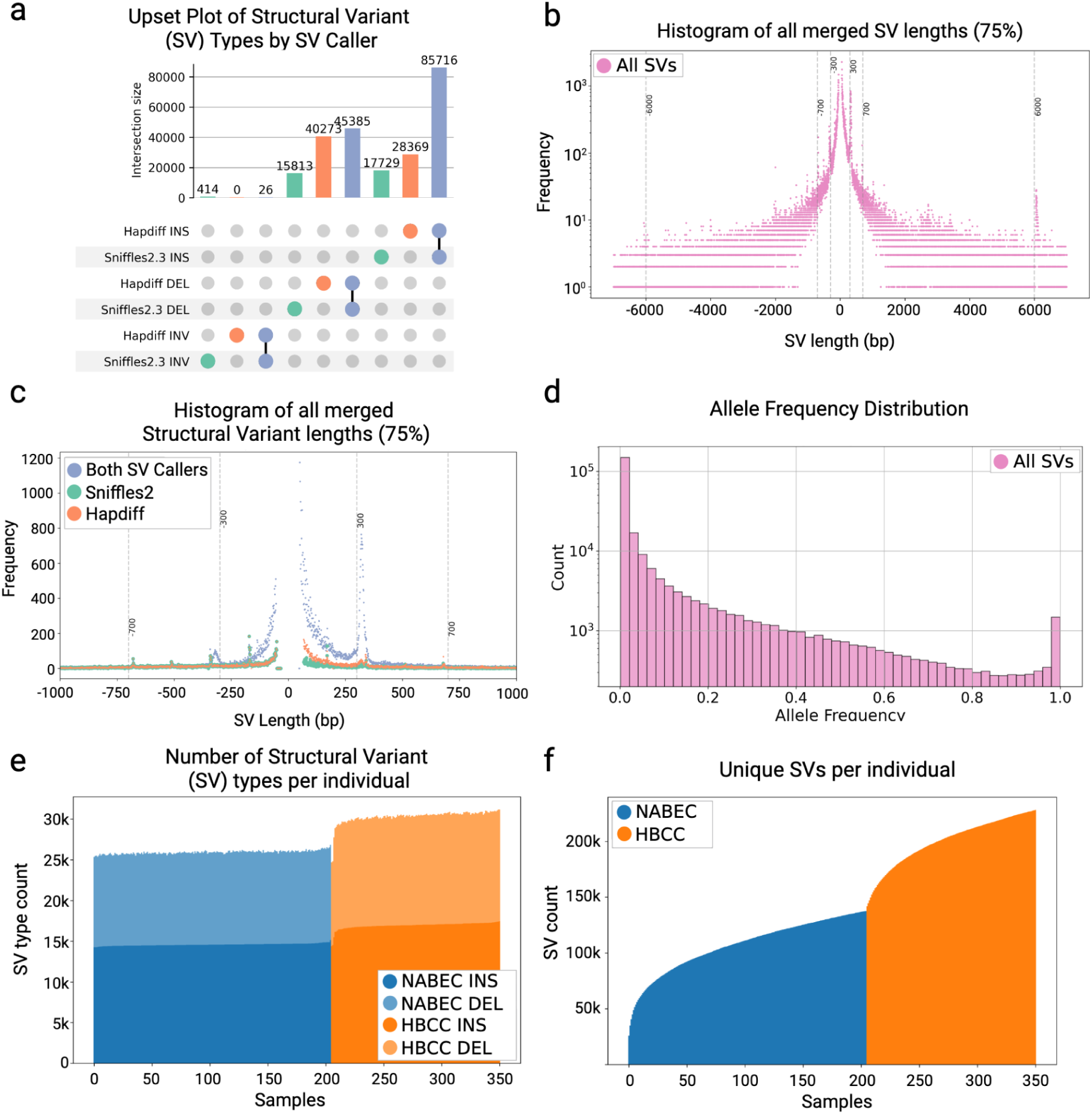
Structural variant characterization across both cohorts. **a.** An upset plot of SVs by SV caller and SV type tallied after merging. Hapdiff called 28,369 insertions and 40,273 insertions that weren’t merged with Sniffles. Sniffles called 17,729 insertions and 15,813 deletions that weren’t merged with Hapdiff. 85,716 insertions and 45,385 deletions were called by both callers and merged together by Truvari. **b.** The size of insertions and deletions is plotted as negative for deletions and positive for insertions. Expected peaks of common SVs are shown at 300 bps, Alu repeats, 700 bps, SINE, and 6000 bps LINE1. **c.** Size of SVs that are merged between SV callers, shown in lavender, and called by just one SV caller. Sniffles SVs that weren’t merged with Hapdiff SVs are shown in green and Hapdiff only SVs are in orange. **d.** Allele frequency distribution of merged SVs. **e.** The number of SVs per individual is stratified by insertions in dark blue, NABEC, or dark orange, HBCC, and deletions in the lighter blue and orange. **f.** A cumulative bar chart that shows the number of new unique SVs added by each individual across both cohorts. The European NABEC cohort contributed ∼115,000 SVs and the HBCC cohort contributed another ∼125,000 SVs to create the set of 234,905 SVs merged across both cohorts and SV callers.

In the NABEC cohort, we detected on average 25,937 SVs per genome (Figure 2e) which impacted an average 15.4 Mb of inserted genomic sequences and 7.4 Mb of deleted sequence per individual. As expected given its ancestry^24^ the HBCC cohort yielded more SVs per genome, averaging 30,322 high-confidence SVs that impacted 15.7 Mb of inserted sequence and 7.9 Mb of deleted sequence per individual (Figure 2e). Most SVs per sample were insertions (average 15,556; 55.98%) and deletions (12,204; 43.89%), with inversions less frequent (31 inversions on average (0.12%)). These results are consistent with findings from other large-scale long-read sequencing studies, such as the 1000 Genomes Project^25^, which reported an average of 24,543 SVs per genome, the Human Genome Structural Variation Consortium (HGSVC^24^), which reported an average of 27,622 SVs per genome, the DECODE Icelandic study with 22,636 SVs per genome^26^, and the NIH All of Us study, which identified 24,285 SVs per genome^27^.

Across both cohorts, we identified 234,905 high-confidence SVs, consisting of 131,813 insertions, 102,651 deletions, and 440 inversions. Among the 90,580 SVs that intersect protein-coding genes, 93.1% (84,296) were located in intronic regions, 6.9% (6,234) overlapped coding regions, and 4.1% (3,715) were found within or spanned the 5’ or 3’ untranslated regions. The average SV size was 3,681 bp, ranging from 50 bp to 76 Kbp. As observed in previous studies, insertion sizes peaked at approximately 300 bp and 6 kb, corresponding to *Alu* and LINE1 transposable elements, respectively (Figure 2b) ^25,28^. We annotated the SVs with known clinical SV databases (nstd102 in dbVar) or any DGV SVs (GRCh38_hg38_variants_2020-02-25.txt) using sveval^29,30^ and, as expected, no known pathogenic SVs were detected in these control samples. We next assessed the population allele frequencies of SVs identified in our study by comparing them to frequencies reported in 1,108 long-read genomes from the 1000 Genomes Project and the Human Pangenome Reference Consortium (HPRC) using STIX^31^. A significant linear correlation was observed when comparing allele frequencies from NABEC to EUR STIX frequencies and HBCC to AFR STIX frequencies, indicating strong concordance between our data and these reference datasets (Supplementary Figure 2). Using the STIX annotations, we estimate that 32.5% of SVs across the two cohorts were novel, defined as variants not reported in previous large-scale long-read sequencing studies.

### The impact of structural variants on gene expression in the human frontal cortex

We performed an SV-only eQTL analysis to evaluate the overall impact of SVs on gene expression in the human frontal cortex. Focusing on common variants (MAF > 5% within this cohort), we investigated their cis-regulatory effects using gene-level whole-transcriptome data from short-read RNA-seq.^32–34^ Expression data were available for 205 of the NABEC samples and 76 of the HBCC samples^35^. After filtering, 24,375 SVs were included in the NABEC cohort expression quantitative trait loci (eQTL) analysis along with expression data for 25,887 expressed genes, of which 15,502 were protein-coding. In the HBCC cohort, 38,018 SVs were used in the eQTL analysis with expression data for 25,847 expressed genes, 15,401 of those being protein-coding genes. An eQTL was defined as an eVariant/eGene pair for all variant/gene pairs within a *cis* window that included variants located within 1 Mb of the gene’s transcriptional start site (TSS).

Using TensorQTL and applying a standard false discovery rate (FDR) threshold of q-value < 0.05, we identified 838 significant SV-eQTL (q-value <0.05) in the NABEC datasets affecting 779 distinct eGenes and 135 significant SV-eQTL in the HBCC datasets affecting 134 distinct eGenes (Table 1, Supplemental Figure 6). 64 of the significant SV-eQTL hits were common across both cohorts (NABEC: HBCC; 8.2%: 54.2%) and 43 SV-eQTLs had the same SV as the top hit per gene. To assess SV genotyping accuracy, we manually reviewed 30 randomly selected significant SV-eQTL hits in IGV, verifying each call by inspecting read coverage changes around breakpoints in the corresponding BAM files. We were able to validate 26/30 NABEC SVs (86.7%) and 23/30 (76.7%) HBCC SVs. In non-coding SV-eQTLs the effect directions (β) were bimodal across SV types, reflecting both increased and decreased gene expression. Consistent with previous studies, exon-overlapping deletion SVs were generally associated with decreased gene expression^36^ (Supplementary Figure 3). Similarly, in line with short-read-based SV-QTL studies, we observed that many insertion exon-overlapping SV-eQTLs resulted in increased gene expression (Supplementary Figure 3).

**Table 1:**
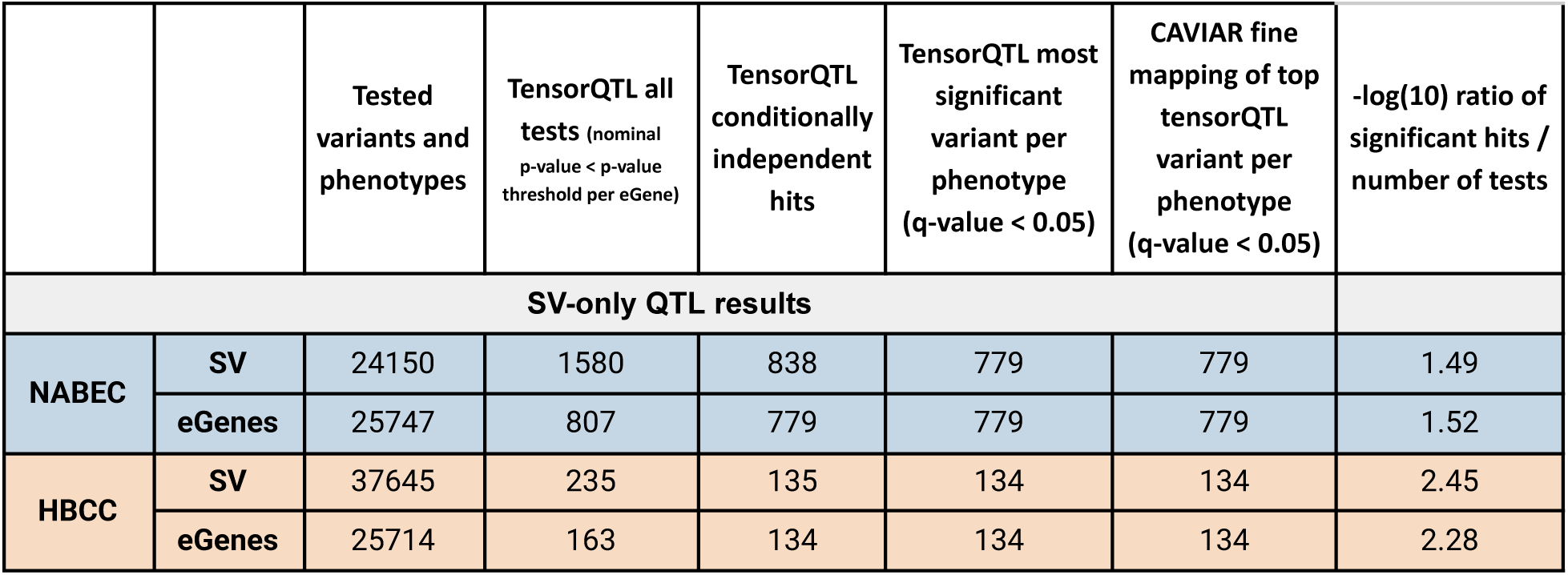
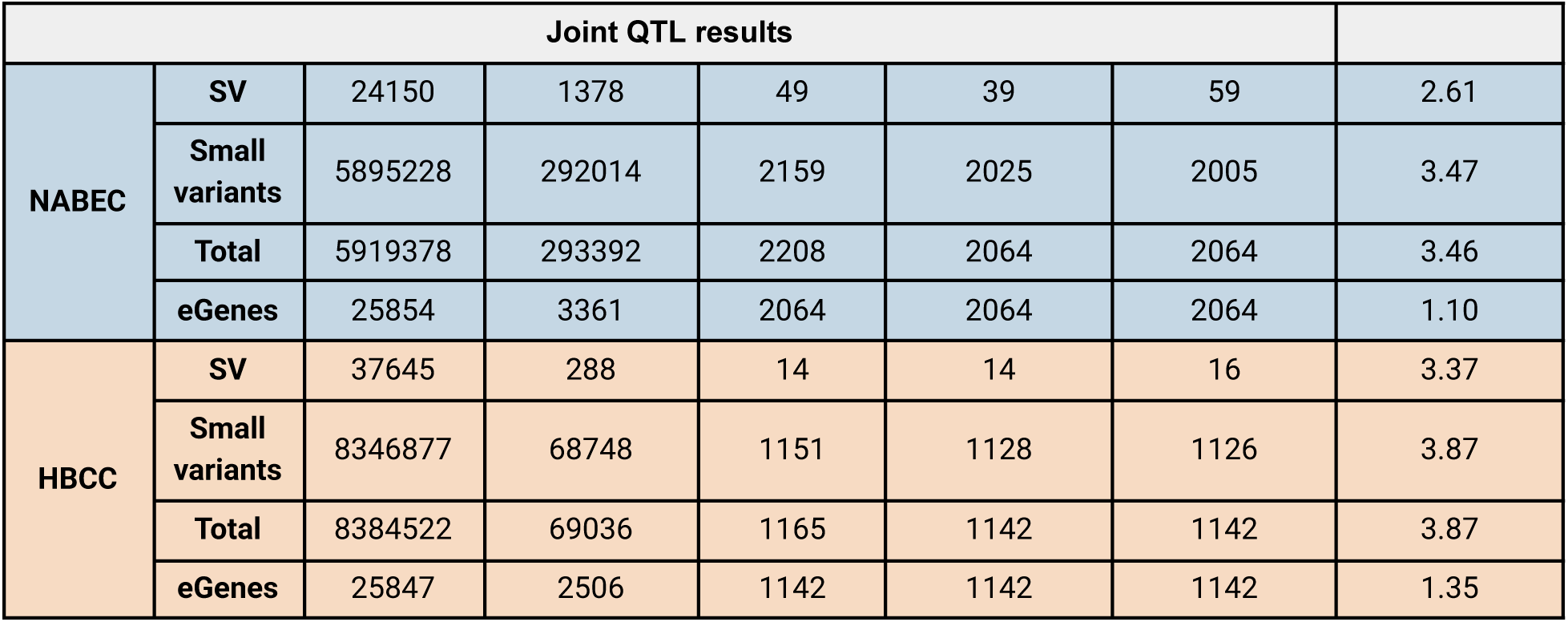
The number of eQTLs in SV only and SV+SNV tensorQTL analysis stratified by cohort and variant type. Significant hits counts are in the TensorQTL conditionally independent hits column and the counts for most significant variant per gene are in the TensorQTL most significant variant per phenotype column.

|  |  | Tested<br>variants and<br>phenotypes | TensorQTL all<br>tests (nominal<br>p-value < p-value<br>threshold per eGene) | TensorQTL<br>conditionally<br>independent<br>hits | TensorQTL most<br>significant<br>variant per<br>phenotype<br>(q-value < 0.05) | CAVIAR fine<br>mapping of top<br>tensorQTL<br>variant per<br>phenotype<br>(q-value < 0.05) | -log(10) ratio of<br>significant hits /<br>number of tests |
| --- | --- | --- | --- | --- | --- | --- | --- |
| SV-only QTL results |  |  |  |  |  |  |  |
| NABEC | SV | 24150 | 1580 | 838 | 779 | 779 | 1.49 |
|  | eGenes | 25747 | 807 | 779 | 779 | 779 | 1.52 |
| HBCC | SV | 37645 | 235 | 135 | 134 | 134 | 2.45 |
|  | eGenes | 25714 | 163 | 134 | 134 | 134 | 2.28 |

**Table 1:** The number of eQTLs in SV only and SV+SNV tensorQTL analysis stratified by cohort and variant type.
| Joint QTL results |  |  |  |  |  |  |  |
| --- | --- | --- | --- | --- | --- | --- | --- |
| NABEC | SV | 24150 | 1378 | 49 | 39 | 59 | 2.61 |
|  | Small variants | 5895228 | 292014 | 2159 | 2025 | 2005 | 3.47 |
|  | Total | 5919378 | 293392 | 2208 | 2064 | 2064 | 3.46 |
|  | eGenes | 25854 | 3361 | 2064 | 2064 | 2064 | 1.10 |
| HBCC | SV | 37645 | 288 | 14 | 14 | 16 | 3.37 |
|  | Small variants | 8346877 | 68748 | 1151 | 1128 | 1126 | 3.87 |
|  | Total | 8384522 | 69036 | 1165 | 1142 | 1142 | 3.87 |
|  | eGenes | 25847 | 2506 | 1142 | 1142 | 1142 | 1.35 |

To further assess the contribution of SVs to frontal cortex gene expression, we performed a joint eQTL analysis combining SVs and small variants (SNVs and indels). Due to the lower SNV calling accuracy of ONT R9 compared to ONT R10, we used Illumina-based SNV data for NABEC and ONT-based SNV data for HBCC. In this SV-SNV eQTL we observed a 2.6 and 8.6 fold increase in eQTLs for NABEC and HBCC after including the 5 and 8 million SNVs for each of these cohorts. In the NABEC joint SV and SNV eQTL analysis, we identified 2,208 significant eQTLs affecting 2,064 distinct eGenes, 49 of which had an SV as the lead variant. In HBCC, we identified 1,165 eQTLs affecting 1,142 distinct eGenes, 14 of which had an SV as the lead variant (Table 1). Notably, 563 eQTLs were shared between the cohorts (NABEC: HBCC; 27.3%: 50.4%), which aligns with previous SNV-based QTL studies reporting approximately 30% overlap in eQTL across cohorts of European and African admixed ancestry^37^. Still, the amount of overlap may be influenced by the difference in samples with RNA-seq data between the NABEC and HBCC cohorts.

### SVs as top candidate variants driving eQTL associations in the human frontal cortex

To assess the potential causal role of SVs at each locus, we performed statistical fine-mapping using Causal Variants Identification in Associated Regions (CAVIAR), estimating the posterior probability of each SV being causal compared to nearby small variants within a 1 Mbp window, while accounting for linkage disequilibrium (LD) between variants^38^. CAVIAR identified an SV as the most likely causal variant in 2.9% (59/2064) of NABEC eQTLs and 1.4% (16/1142) of HBCC eQTLs (Table 1). These results align with findings from Kirsche et al., who also applied long-read-derived SV-eQTL fine-mapping and reported high posterior probabilities for SVs in 3.65% of eGenes^39^. Although SV eQTL counts are lower than SNV eQTLs here, SVs are more likely to be an eQTL on a per-variant basis. In the NABEC cohort 0.2% (59/24150) of SVs were top QTL hits while 0.03% of tested SNVs ended up being QTL hits and similarly in the HBCC cohort 0.04% (16/37645) of SVs and 0.01% of SNVs were QTL hits. Of the 563 eGenes that were significant SV+SNV eQTLs, 59 NABEC and 16 HBCC eGenes have SVs as the top variant after fine mapping. These SV driven eQTLs are distributed across the genome with effect sizes comparable to or higher than surrounding SNV driven eQTLs (Supplementary Figure 6). All of the 59 NABEC SV+SNV eGenes with a lead SV were identified in the SV-eQTL, as were 13/16 HBCC SV+SNV eGenes. GeneOntology of these lead SV eGenes show that they are involved in several biological functions including immune response, DNA binding, structural proteins, and protein modification ^40^. This work suggests that many previously undetected SVs may play a significant role in differential gene expression, compared to the SNVs identified in earlier eQTL studies^39^.

Among the 64 candidate causal SVs identified across both cohorts, one notable example is located at the *NLRP2* locus. *NLRP2*, part of the nucleotide-binding and leucine-rich repeat receptor (NLR) family of immune-related genes, plays a key role in regulating inflammatory responses^41^. A 1,090 bp deletion (napu_chr19_54964613_54965703_DEL_-1090) in the 5’ untranscribed region (UTR) of *NLRP2* is associated with increased gene expression in the frontal cortex within the HBCC cohort (p-value = 1.79x10^-7^; FDR-adjusted q-value = 8.99×10⁻^4^) (Figure 3a). However, in the NABEC cohort, this deletion was excluded from the QTL analysis due to a low genotyping rate, reflecting SV merging artifacts. Instead, another SV tagging the deletion, a 1,382 bp insertion (napu_chr19_54965640_54965640_INS_1382) was identified as the candidate causal variant. This insertion is in high LD with the deletion (R² = 0.97, D’ = 1) and both are positively associated with *NLRP2* gene expression, in Figure 3a, however, the expression is plotted for the genotypes of the deletion. Another example of an SV eQTL, which was only significant in the NABEC cohort, is a 172 bp insertion (napu_chr13_111326259_111326259_INS_172) located within the gene *TEX29* (Testis Expressed 29). This insertion was associated with increased gene expression in carriers (NABEC p-value = 5.96×10^-17^; q-value = 7.74×10^-11^) (Figure 3b). At the time of this work TEX29 expression in the brain has not previously been reported. In the HBCC cohort, increased expression of the TEX29 gene is associated with a SNP (chr13:111329944:C:T).

**Figure 3:**
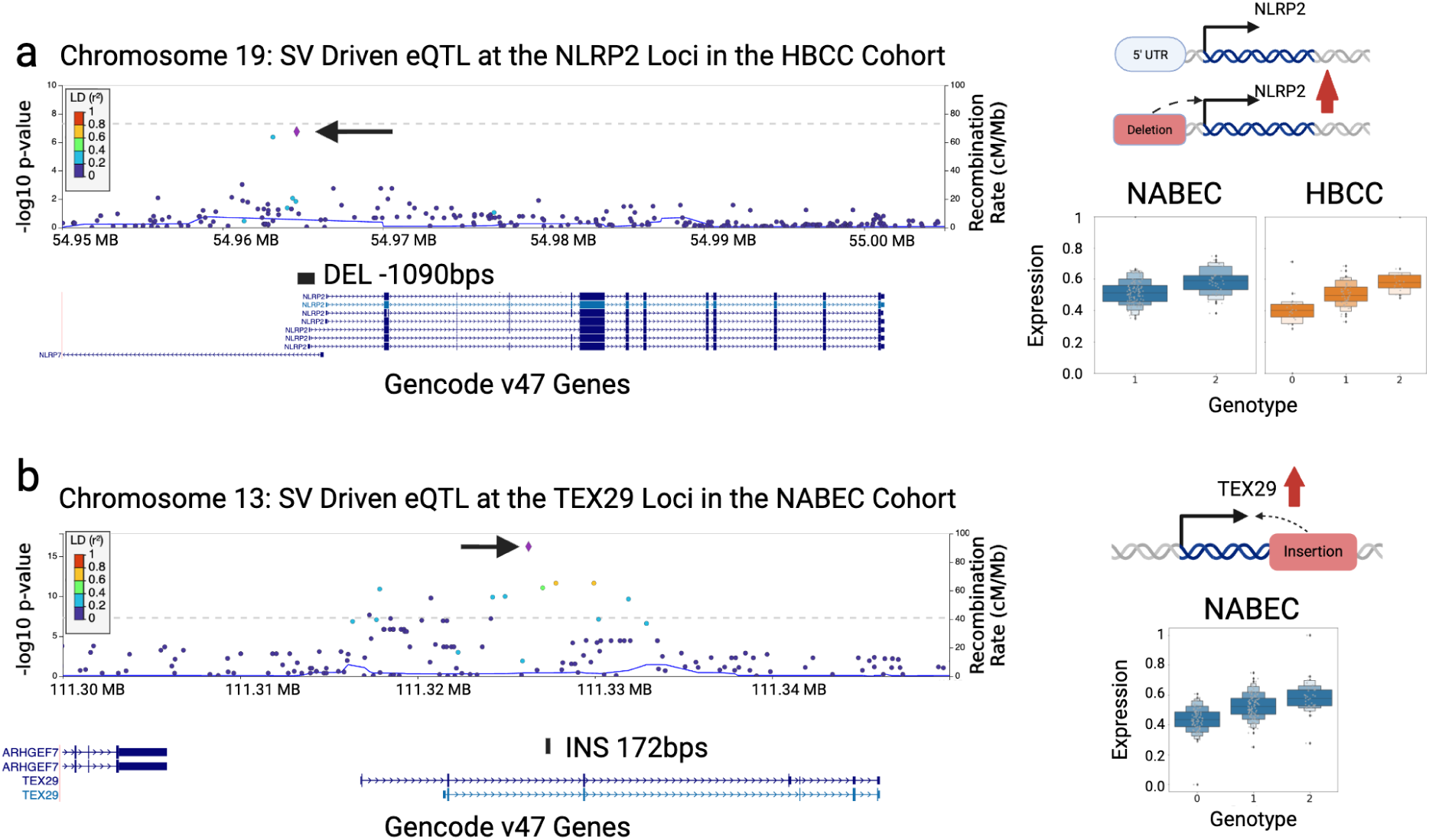
eQTL discovery and fine-mapping. **a.** Example of an eQTL in the NLRP2 gene. The top panel presents a locuszoom plot of the SV and small variant joint eQTL for HBCC, with an arrow indicating the top significant variant, napu_chr19_54964613_54965703_DEL_-1090. Underneath is a depiction of a deletion overlapping the 5’ transcription start sites of several transcripts of NLRP2. On right are boxplots of the eQTL stratified by genotypes of the deletion (NABEC in blue, HBCC in orange). The top right panel depicts a schematic representation of the effect of the variant. **b.** Example of an SV and small variant joint eQTL driven by an SV in NABEC. The top panel presents a locus zoom plot of the SV and small variant joint eQTL for NABEC, with an arrow indicating the top significant variant, napu_chr13_111326259_111326259_INS_172. Underneath the variant is depicted overlapping an intron of *TEX29*. The right panel displays a boxplot of the eQTL stratified by genotypes of the insertion (NABEC in blue). The top right panel depicts a schematic representation of the variant’s effect.

### Genome-wide Methylation profiling in the human frontal cortex

LRS enables the simultaneous detection of both sequence variation and DNA methylation directly from native DNA. LRS-based methods provide more complete and contiguous methylation profiles than short-read technologies, which often rely on indirect methods like bisulfite conversion and can miss key regulatory regions due to fragmented or incomplete coverage^42^. Prior work has shown that methylation calls in these brain tissue samples reflect the expected distribution of methylation patterns for primary tissue, and in this case prefrontal cortex tissue specifically^43^. The majority of CpG’s are hypermethylated (>60% methylation frequency) in these tissues and less than 10% of whole genome windows are hypomethylated (<60% methylation frequency). Methylation was aggregated across the genome by averaging over windows of at least 1 Kbp and expanding the window to include a minimum of 50 CpGs. The resulting sizes of these windows range from 1Kb to over 10Kbp, although only 10% are 10Kbp or greater. The technical differences in ONT R9 and R10 methylation measurements, shown in previous work to be more precise in R10 data, are apparent in Figure 4a (Supplementary Figure 4)^44^. Aggregating methylation calls, however, can minimize this difference. Due to this difference, though, we analyzed the samples in the NABEC and HBCC cohorts separately.

**Figure 4:**
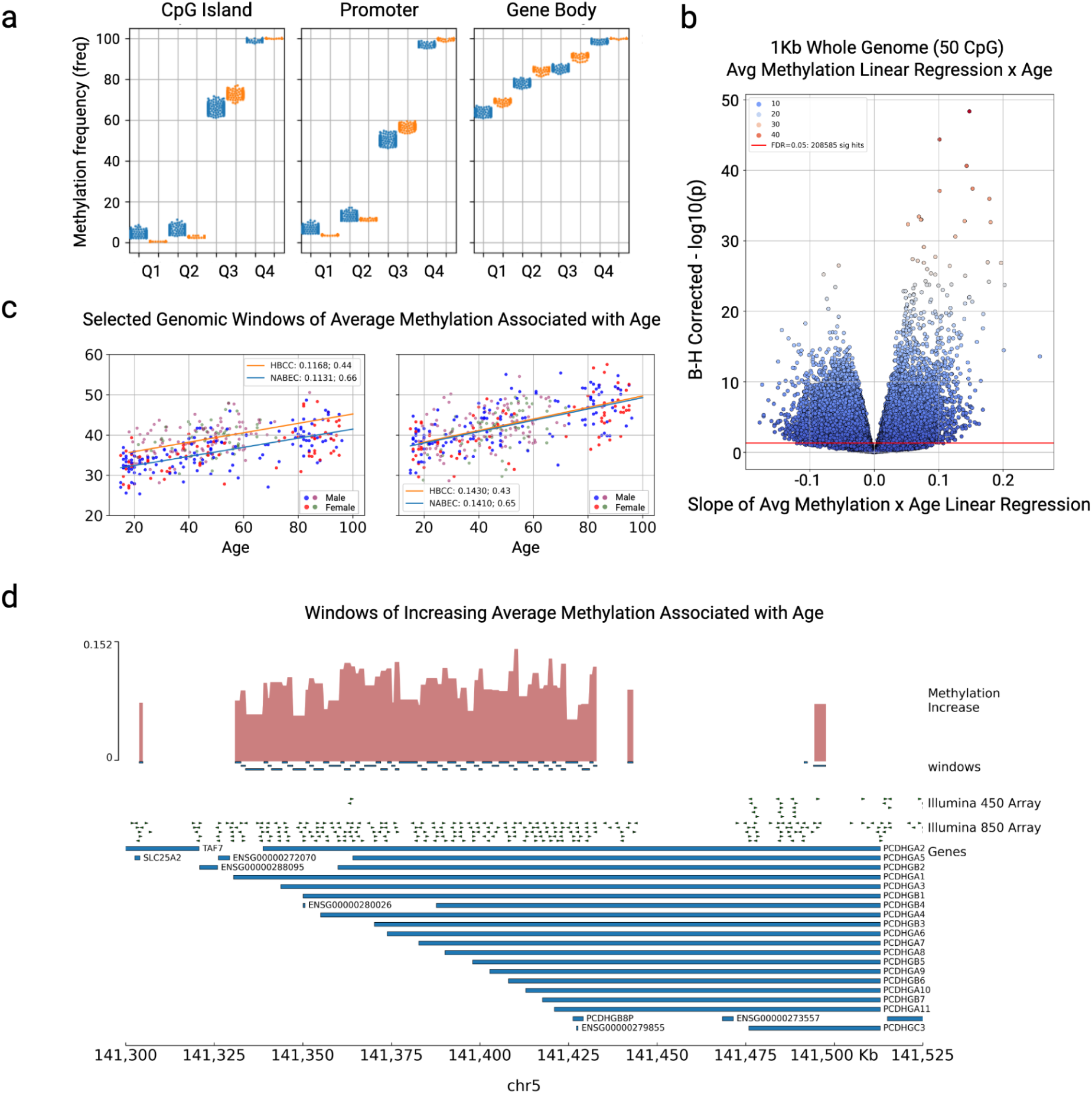
Profiling genome-wide methylation in the frontal cortex. **a.** Aggregated methylation frequency of NABEC (blue) and HBCC (orange) samples by quartile for CpG islands, promoter regions (extended to 2kb), and across RefSeq gene bodies. **b.** Age associations of whole genome 1Kb / 50 CpG minimum windows of averaged methylation in the NABEC cohort. Many regions (145,086; FDR 0.05) were weakly associated with age; the slope of the linear regression ranged from -0.1 to 0.1. The volcano plot is of age regressions; x-axis is the slope of the regression line y-axis is the Benjamini-Hochberg corrected p-values. **c.** Two neighboring regions significantly associated with age that overlap exons of the Protocadherins Gamma (PCDHAG) cluster of genes. **d.** Tracks of windows showing methylation increase with age. Top track is a barplot of the amount of methylation increase (slope) by age. Window locations associated with age are shown below. The locations of Illumina 450 and EPIC 850 Methylation Arrays are shown above the gene locations. Across the bottom are coordinates of Chr 5.

Using the 1Kb windows and CpG islands (CGIs) we looked for regions whose methylation status is correlated with age. To do this we performed an ordinary least squares linear regression with age as the independent variable and average methylation frequency as the dependent variable. It is important to note that the NABEC cohort has 51 more individuals as well as longer-lived individuals on average than the HBCC cohort and thus had stronger correlations with age. Moreover, the admixture of the African American samples may contribute to the smaller number of correlations with age^45^. Around 28% of the 518,000 whole genome windows and 14% of the CGIs in the NABEC cohort were significantly (FDR < 0.05) associated with age. Similarly in HBCC 8.5% of the 518,466 coverage filtered windows and 20% of the CGIs in the HBCC cohort were significantly (FDR < 0.05) associated with age (Figure 4b). Across cohorts there was agreement in 5% of the genome windows and 1% of the CGIs showing significance in both cohorts and in the same direction (Supplementary Table S21). For most CGIs and genomic windows passing multiple test correction, the change in methylation represented by the regression slope was subtle (less than 0.1). However, in CGIs and 1Kb windows with the largest methylation changes (slopes greater than 0.1 and an R^2^ of more than 0.4) they do overlap with some functionally relevant regions. For example, many regions significantly associated with age overlap the protocadherin alpha and gamma clusters of genes on the q-arm of chromosome 5 (Figure 4c, 4d). Genes from these clusters create protein products that combine to produce unique protein markers on the outside of each neuron in the human brain responsible for distinguishing a neighboring neuron from itself.^46^ Fewer regions showed a decrease in methylation with age. Regions with the largest decrease in methylation frequency by age were found within the *FKBP5* gene, which functions as a mediator of glucocorticoid receptor cortisol binding and signaling^47^. The *FKBP5* protein product, FKBP51, binds to glucocorticoid receptors and is mediated by its epigenetic status and ultimately can prolong the body’s stress response through its impact on the levels of circulating cortisol and glucocorticoids^47,48^. Interestingly, overexpression of *FKBP5* may also play a role in *Tau* protein accumulation which is linked to AD^48^.

### Structural variants influence DNA methylation in the human frontal cortex

Genetic variation plays a critical role in shaping DNA methylation patterns in the human brain, influencing gene regulation and contributing to neurodevelopment and disease complexity ^49,50^. While small variants have been well studied in this context, the influence of SVs remains less understood due to challenges in detection and genotyping. Building on the eQTL framework, we performed mQTL analyzes to evaluate the effects of SVs on DNA methylation in the postmortem frontal cortex, focusing on CGIs (defined as the 27,949 CpG Islands in GRCh38, having more than twice the expected number of CG dinucleotides based on CG content of the rest of the genome), promoter regions, and gene bodies. ^51^. We defined an mQTL as cis-acting variants that significantly influence DNA methylation within a 1 Mb window surrounding each CGI, promoter, or gene body. Hence in this context, cis-acting refers specifically to variants affecting any methylation within this 1 Mb range. Hereafter, we refer to these as CGI-mQTL, prom-mQTL, and gene-mQTL, respectively. We ran these mQTLs in two ways first using all CpGs present in any sample, then to maintain a consistent number of CpGs across samples we removed CpGs in all samples if they were within the boundaries of an SV deletion in any sample. This is particularly important for deletions that overlap the methylation region of interest causing a variable number of CpGs to be aggregated in samples that carry the deletion compared to those that don’t. We present results from both approaches, however by removing deletions we can be certain that the methylation associated with genetic variation is not confounded by the presence or absence of CpG sites in a subset of samples.

We first conducted an SV-only mQTL analysis with tensorQTL identifying 4,151 CGI, promoter, or gene body mQTLs across both cohorts. Similar to the SV-eQTL analysis, we observed a bimodal distribution of effect directions (β) between SV types for non-coding SV-mQTLs at CGIs, promoters, and gene bodies. Two thirds of the significant hits were deletions, 4 of which removed entire CpG Islands, and 57% of SVs were identified within the methylation region of interest with variable effect sizes. When we removed deleted CpGs from this analysis, 1,385 SV-only mQTLs remained significant across regions and cohorts. Notably, when focusing on SV-mQTLs overlapping exons, SV-prom-mQTLs displayed a distinct pattern: insertions were generally associated with hypermethylation, while deletions were linked to hypomethylation (Supplementary Figure 5).

In the NABEC cohort, we identified 612 SV-CGI-mQTLs across 567 distinct CGIs. Similarly, in the HBCC cohort 266 SV-CGI-mQTL were identified across 259 distinct CGIs. Of these, 100 (NABEC: HBCC; 17.6%: 38.6%) were shared between both cohorts, exhibiting the same phenotypes and consistent directions of effect (β). Next, we examined promoter regions, identifying 652 SV-prom-mQTL in NABEC across 614 distinct promoters and 486 SV-prom-mQTL across 472 distinct promoters in HBCC (Table 2). A total of 117 (NABEC: HBCC; 19.1%: 24.9%) SV-prom-mQTLs were common across both cohorts, with the same SVs and direction of effect (β). The overlap observed here is smaller than that seen between eQTL hits across both cohorts, and may be due to greater variability in the methylation data compared to the expression data, as NABEC samples were sequenced on R9, while HBCC samples were sequenced on R10. For gene bodies, we identified 1687 SV-gene-mQTL in NABEC for 1581 distinct genes and 682 SV-gene-mQTL in HBCC for 658 distinct genes (Table 2), with 252 (NABEC: HBCC; 16.0%, 37.0%) common between both cohorts, showing consistent SVs and effects (β).

**Table 2:** The number of tensorQTL mQTLs in SV only and SV+SNV tensorQTL analysis stratified by cohort and variant type. The number of SVs or SNVs is reported by tensorQTL and after CAVIAR fine-mapping selecting the most likely variant.

|  |  | Tested<br>variants and<br>phenotypes | tensorQTL all<br>tests (nominal<br>p-value <<br>p-value<br>threshold per<br>eGene) | tensorQTL<br>conditionally<br>independent<br>hits | tensorQTL<br>most<br>significant<br>variant per<br>phenotype<br>(q-value <<br>0.05) | tensorQTL<br>most<br>significant<br>variant per<br>phenotype<br>(q-value <<br>0.05)<br>delRemoved | CAVIAR fine<br>mapping of<br>top<br>tensorQTL<br>variant per<br>phenotype<br>(q-value <<br>0.05) |
| --- | --- | --- | --- | --- | --- | --- | --- |
| SV meth QTL results |  |  |  |  |  |  |  |
| NABEC |  | CGI sites |  |  |  |  |  |
|  | SV | 23681 | 1143 | 612 | 567 | 392 | 567 |
|  | sites | 25472 | 589 | 567 | 567 | 392 | 567 |
|  |  | Promoters |  |  |  |  |  |
|  | SV | 23232 | 1019 | 652 | 614 | 302 | 614 |
|  | sites | 27407 | 658 | 614 | 614 | 302 | 614 |
|  |  | Gene bodies |  |  |  |  |  |
|  | SV | 24351 | 3019 | 1687 | 1581 | 336 | 1581 |
|  | sites | 40990 | 1678 | 1581 | 1581 | 336 | 1581 |
| HBCC |  | CGI sites |  |  |  |  |  |
|  | SV | 34751 | 451 | 266 | 259 | 133 | 259 |
|  | sites | 24984 | 289 | 259 | 259 | 133 | 259 |
|  |  | Promoters |  |  |  |  |  |
|  | SV | 34131 | 646 | 486 | 472 | 139 | 472 |
|  | sites | 27197 | 523 | 472 | 472 | 139 | 472 |
|  |  | Gene bodies |  |  |  |  |  |
|  | SV | 35711 | 1069 | 682 | 658 | 83 | 658 |
|  | sites | 40707 | 733 | 658 | 658 | 83 | 658 |
| Joint meth QTL results |  |  |  |  |  |  |  |
| NABEC |  | CGI sites |  |  |  |  |  |
|  | SV | 23681 | 1092 | 69 | 60 | 43 | 68 |
|  | SNV | 5726688 | 171888 | 1691 | 1563 | 1006 | 1555 |
|  | Total | 5750369 | 172980 | 1760 | 1623 | 1049 | 1623 |
|  | sites | 25603 | 2437 | 1623 | 1623 | 1049 | 1623 |
|  |  | Promoters |  |  |  |  |  |
|  | SV | 23232 | 908 | 36 | 30 | 21 | 56 |
|  | SNV | 5646975 | 211618 | 2121 | 1991 | 875 | 1965 |
|  | Total | 5670207 | 212526 | 2157 | 2021 | 929 | 2021 |
|  | sites | 27422 | 3242 | 2021 | 2021 | 929 | 2021 |
|  |  | Gene bodies |  |  |  |  |  |
|  | SV | 24351 | 2570 | 130 | 120 | 25 | 167 |
|  | SNV | 5943646 | 538688 | 4172 | 3906 | 904 | 3859 |
|  | Total | 5967997 | 541258 | 4302 | 4026 | 896 | 4026 |
|  | sites | 41140 | 6600 | 4026 | 4026 | 896 | 4026 |
| HBCC |  | CGI sites |  |  |  |  |  |
|  | SV | 34751 | 355 | 39 | 36 | 26 | 37 |
|  | SNV | 8194130 | 60784 | 1012 | 976 | 527 | 975 |
|  | Total | 8228881 | 61139 | 1051 | 1012 | 553 | 1012 |
|  | sites | 25131 | 1867 | 1012 | 1012 | 553 | 1012 |
|  |  | Promoters |  |  |  |  |  |
|  | SV | 34131 | 510 | 38 | 37 | 11 | 44 |
|  | SNV | 8055277 | 126480 | 2296 | 2199 | 765 | 2192 |
|  | Total | 8089408 | 126990 | 2334 | 2236 | 776 | 2236 |
|  | sites | 27213 | 3937 | 2236 | 2236 | 776 | 2236 |
|  |  | Gene bodies |  |  |  |  |  |
|  | SV | 35711 | 778 | 54 | 52 | 4 | 78 |
|  | SNV | 8501372 | 162216 | 2042 | 1971 | 376 | 1945 |
|  | Total | 8537083 | 162994 | 2096 | 2023 | 380 | 2023 |
|  | sites | 40876 | 4059 | 2023 | 2023 | 380 | 2023 |

Next to further assess the relative contribution of SVs to methylation in the postmortem frontal cortex, we conducted a joint QTL analysis combining SVs and small variants (Table 2). For the CGI-mQTL, prom-mQTL, and gene-mQTL, 419 (NABEC: HBCC; 25.8%: 41.4%), 676 (NABEC: HBCC; 33.4%: 30.2%), and 809 (NABEC: HBCC; 20.1%: 40.0%) were shared between both cohorts, with the same phenotype and direction of effect (β). A total of 12,941 SV-SNV-mQTLs were originally identified, however, after removing deletions and re-running the joint SV-SNV-mQTLs 4,583 mQTLs remained significant across regions and cohorts. Overall, the odds ratio for an SV to be mQTL compared to small variants was 9.9 (95% CI: 8.16-11.91), 4.3 (95% CI: 3.44-5.48), and 6.5 (95% CI 5.54-7.55) for CGI-mQTL, prom-mQTL, and gene-mQTL, respectively, across both datasets, suggesting that individual SVs are more likely than individual small variants to act as an mQTL (Supplementary Table 4). Interestingly, the first promoter for the TEX29 gene was a SV-SNV-prom-mQTL in both cohorts and the same SNP (chr13:111321006:A:G) was associated with reduced promoter methylation; TEX29 being the same eGene with increased expression associated with an SV in the NABEC cohort (Figure 3). Overall four eGenes with SVs as the lead eQTL variant were also SV-SNV-prom-mQTLs: NLRP2, METTL21C, SMN2, and TEX29. Though only NLRP2 and METTL21C had the same SV as the eQTL and mQTL lead variant, SMN2 and TEX29 had an SNP as the lead mQTL variant. The genes associated with promoters in SV-SNV-prom-mQTLs were found to have biological functions that include neuron projection, brain development, synapse assembly, myelination, cell recognition, DNA methylation, and high density lipoproteins in extracellular space according to their gene ontology annotations.

There were fewer significant mQTLs in the runs with deleted CpGs removed for all 3 regions tested, however, we present both sets of results because some potentially biologically relevant mQTLs were only identified using raw methylation data. For example, the HLA class I histocompatibility antigen, C alpha chain (HLA-C), Synuclein Alpha (SCNA), and the Amyloid-beta A4 precursor protein-binding family B members: APBB3 and APBB1IP are only mQTLs in the raw methylation mQTLs. Except for gene bodies, 30-80% of deletion filtered mQTL phenotypes were also identified in the corresponding raw methylation mQTL. Due to their large size many more CpGs were removed from deletions within gene bodies than other phenotypic regions and caused an 80% drop in gene body SV-mQTLs and SV+SNV-mQTLs. Across all the deletion filtered mQTLs over 1500 new mQTLs were contributed and supplementary mQTL tables contain both sets of results. One new mQTL of note is at the LHX6 promoter in the NABEC SV+SNV-PROM-mQTL where a SNP was associated with a slight decrease in promoter methylation, this gene is involved with tumor suppression, GABAergic interneuron development, and migration and forebrain development ^52,53^.

### SVs are amongst the top candidate variants driving mQTL associations in the human frontal cortex

To further assess the potential causal role of SVs at each locus in the joint SV+SNV mQTLs, we performed statistical fine-mapping using CAVIAR. In the NABEC cohort, SVs were identified as the most likely causal variant in 4.2% (68/1623) of CGI-mQTLs, 2.8% (56/2021) of prom-mQTLs, and 4.2% (167/4026) of gene-mQTLs. In the HBCC cohort, SVs were the top probable causal variant in 3.7% (37/1012) of CGI-mQTLs, 2.0% (44/2236) of prom-mQTLs, and 3.9% (78/2023) of gene-mQTLs (Table 2). Across both cohorts, we identified 105 candidate causal SV-CGI-mQTL, 100 SV-prom-mQTL, and 245 SV-gene-mQTL. These SV driven mQTLs are distributed across the genome with effect sizes often higher than surrounding SNV driven mQTLs (Supplementary Figure 6). Among these 450 candidate causal SV-mQTL identified across both cohorts, one notable example is also a SV-eQTL located at the NLRP2 locus. Interestingly the same 1090 bp deletion overlapping the 5’ UTR of the *NLRP2* gene, which was a eQTL hit (napu_chr19_54964613_54965703_DEL_-1090), was also the top candidate causal variant at this locus (Figure 5a). This deletion is significantly associated with frontal cortex hypermethylation of a CGI (chr19_54982861_54983167_CGI_30) that overlaps with exon 6 of NLRP2 in the HBCC cohort (p-value = 2.50x10^-9^; q-value 1.27x10^-3^). This observation is notable because hypermethylation is generally expected to correlate with transcriptional repression. However, in this case, the data suggest a more nuanced regulatory mechanism as this variant was positively associated with gene expression. This SV was a QTL hit after deleted CpGs were removed from all samples as well, which was to be expected as the deletion does not overlap the CGI. Additionally, we identified a 62 bp deletion (napu_chr22_39101634_39101696_DEL_-62) within an intron of *APOBEC3H* as the top candidate causal variant at the *APOBEC3H* locus. *APOBEC3H*, a member of the *APOBEC3* family of cytidine deaminases, plays a crucial role in innate immunity by restricting viral replication, particularly retroviruses, through DNA deamination. This 62 bp deletion is associated (p-value = 1.50x10^-16^; q-value = 7.09x10^-10^) with hypomethylation of an alternative promoter (chr22_39099320_39101380_APOBEC3H_1) in the frontal cortex, further supporting its role as a candidate causal variant (Figure 5b). This example also remained significant in the deleted CpGs removed mQTL analysis in both cohorts.

**Figure 5:**
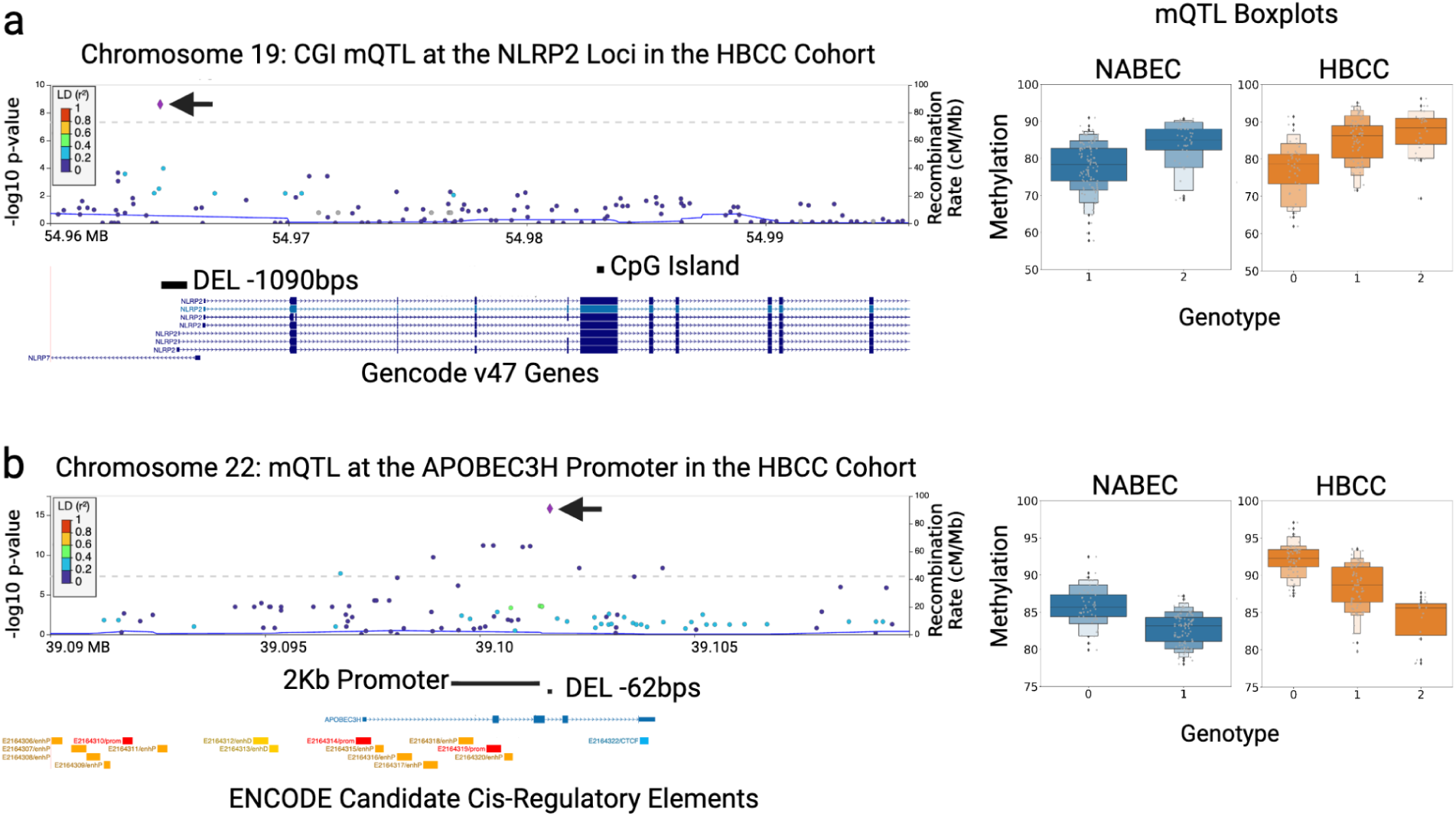
mQTL discovery and fine-mapping. a. Example of a SV-SNV joint cpg-islands-mQTL led by a SV in the gene *NLRP2*. The top panel presents a locuszoom plot of the SV-SNV joint cpg-islands mQTL for HBCC, with an arrow indicating the top significant variant, napu_chr19_54964613_54965703_DEL_-1090. Underneath is a depiction of the variant overlapping the 5’ transcription start sites of several transcripts of NLRP2 and the phenotype cpg island overlapping with the exon of *NLRP2*. The righthand panel shows a boxplot of the eQTL stratified by genotypes of the deletion (NABEC in blue, HBCC in orange). b. Example of a SV-SNV joint promoter-mQTL led by a SV in the gene *APOBEC3H*. The top panel presents a locuszoom plot of the SV-SNV joint cpg-islands mQTL for HBCC, with an arrow indicating the top significant variant, napu_chr22_39101634_39101696_DEL_-62. Below the variant is shown to be overlapping the intron of *APOBEC3H* and the phenotype cpg island overlapping with the exons and intron of *APOBEC3H*. The righthand panel shows a boxplot of the mQTL stratified by genotypes of the deletion (NABEC in blue, HBCC in orange).

### Haplotype-resolved mQTL analysis identifies novel regulatory variants masked in traditional models

Traditional mQTL analysis averages methylation signals across alleles which can obscure allele-specific effects. To address this limitation, we developed ASM-LR, a computational framework for haplotype-resolved mQTL mapping using LRS. For ONT data, methylation is called on a per-read basis by the basecaller and then modkit is used to compute the fraction of modified and canonical bases at each CpG site. When reads are phased these methylation calls can be phased as well. The NAPU pipeline integrates Margin phase for the harmonization of variant genotypes and methylation haplotypes within each sample. Merging and joint genotyping SVs across cohorts, however, remains challenging so the labelling of haplotypes here is arbitrary and assigned locally. ASM-LR performs a linear regression across both haplotypes with cis SVs and SNVs, capturing allele-specific methylation effects irrespective of haplotype labeling. Since ASM-LR uses individual alleles, heterozygous variants are no longer considered in the mQTL calculation thus reducing the noise of genotype groups. Additionally, the number of observations is effectively doubled as two alleles are considered for each individual instead of one. Ideally, the reduced noise and increased observations would increase the statistical power of a phased ASM-LR mQTL compared to an unphased mQTL.

For this study, we focused on a curated set of 171 AD and PD genes ^22,23^ and used ASM-LR to test variants and regions of interest within ±1 Mb of each gene using both raw and deletion filtered methylation data. Across all loci evaluated by ASM-LR, we identified 234 ASM-LR mQTLs across 2,561 CGIs, promoters, and gene bodies in NABEC, while in HBCC, 217 mQTLs were detected across 5,668 regions (Table 3). SVs were the top hit for 62.5% of NABEC ASM-LR mQTLs and 58.4% of HBCC ASM-LR mQTLs and notably SVs were 10 times as likely to be a phased mQTL hit than SNVs in both cohorts. We also ran ASM-LR with deletion filtered methylation data and identified just 64 total ASM-LR mQTLs across both cohorts, 54 NABEC and 10 HBCC mQTLs. Here again we present both deletion filtered and raw methylation because some potentially biologically relevant mQTLs, like a deleted CpG island within a gene body, were only identified using raw methylation data (Figure 6). Though the overall counts were lower for deletion filtered runs, SVs were again the top hit for 65% of NABEC and 57% of HBCC ASM-LR mQTLs. Further, phased models consistently yielded larger effect sizes when compared to unphased mQTLs, underscoring the advantage of haplotype-resolved approaches.

**Figure 6:**
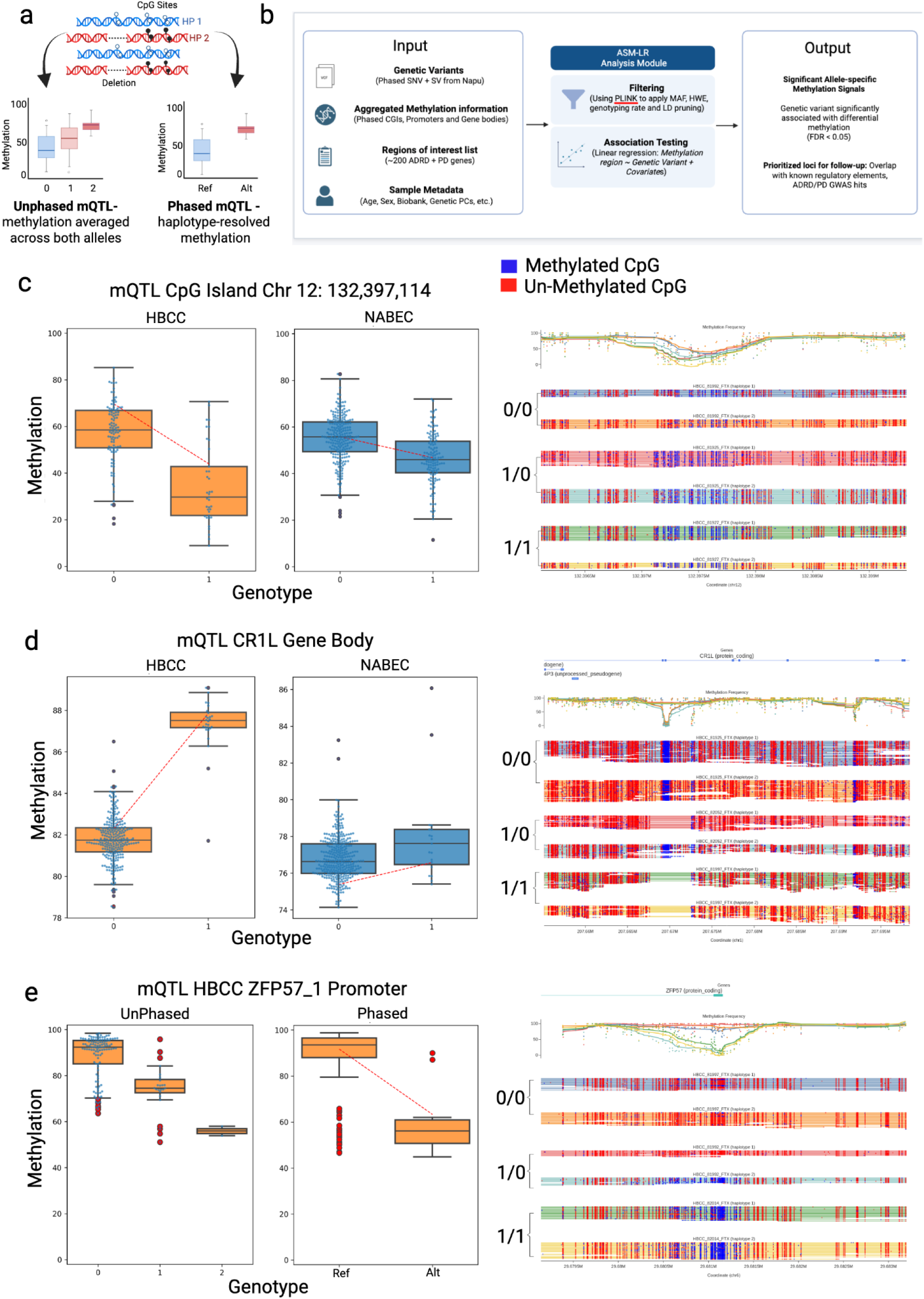
Haplotype-resolved mQTL methods and discovery. a. A depiction of phased reads (haplotype 1 in blue and haplotype 2 in red) with a heterozygous deletion and allele specific methylation patterns. Left-hand boxplot illustrating a unphased mQTL hit where methylation is averaged over both alleles and variant genotypes are converged into dosage genotypes. Right-hand boxplot shows a phased mQTL where methylation is quantified per haplotype the blue reference allele appears hypomethylated and the red alternate allele is, in comparison, hypermethylated. b. Graphical illustration of the ASM-LR haplotype-resolved mQTL framework. c. A phased only mQTL hit with a large methylation change in a CpG Island on chromosome 12 where samples with an insertion is associated with less methylation across the reference CpG positions within the CGI. HBCC cohort is show in orange on the left and the NABEC cohort is show in blue on the right. d. A phased only mQTL hit with a slight methylation increase in alleles carrying a 4.7Kb deletion within the CR1L gene body. e. mQTL hit at the ZFP57_1 promoter region that was identified in both the unphased tensorQTL and phased ASM-LR QTL analysis. A 227 bp deletion was associated with a ∼30% reduction in methylation at the promoter region. All mQTL examples are accompanied by a modbamtools^58^ illustration of the methylation frequency of all mQTL considered CpG positions in phased reads in a homozygous reference, heterozygous, and homozygous alternate sample in the HBCC cohort. Refseq gene tracks are included where applicable.

**Table 3:** The number of tests in each ASM-LR haplotype-resolved mQTL stratified by cohort, region, and variant type. Though more variants were included in the HBCC cohort mQTLs, due to ancestry and sequencing technology, proportionally SVs were more likely to be a leading mQTL variant than an SNV across both cohorts.

|  |  | (n)Tested<br>variant/region<br>pairs | ASM-LR<br>(q-value <<br>0.05) | ASM-LR ratio of<br>significant hits /<br>number of tests | (n)Tested<br>variant/<br>region pairs<br>Dels<br>Removed | ASM-LR<br>(q-value <<br>0.05) Dels<br>Removed | ratio of<br>significant<br>hits /<br>number of<br>tests Del<br>Removed |
| --- | --- | --- | --- | --- | --- | --- | --- |
|  | **ASM-LR phased meth QTL results** |  |  |  |  |  |  |
| **NABEC** |  | **CGI sites** |  |  |  |  |  |
|  | **SV** | 21556 | 27 | 0.001253 | 9531 | 18 | 0.001889 |
|  | **SNV** | 62141 | 16 | 0.000257 | 33797 | 10 | 0.000296 |
|  | **Total** | 83697 | 72 | 0.000860 | 33924 | 28 | 0.000825 |
|  | **sites** | 856 | 43 | 0.050234 | 1443 | 14 | 0.009702 |
|  |  | **Promoters** |  |  |  |  |  |
|  | **SV** | 12160 | 23 | 0.001891 | 3527 | 4 | 0.001134 |
|  | **SNV** | 46273 | 10 | 0.000216 | 20802 | 0 | 0.000000 |
|  | **Total** | 58433 | 38 | 0.000650 | 22081 | 4 | 0.000181 |
|  | **sites** | 658 | 33 | 0.050152 | 707 | 4 | 0.005658 |
|  |  | **Gene bodies** |  |  |  |  |  |
|  | **SV** | 16420 | 50 | 0.003045 | 5133 | 12 | 0.002338 |
|  | **SNV** | 69537 | 34 | 0.000489 | 31055 | 10 | 0.000322 |
|  | **Total** | 85957 | 124 | 0.001443 | 33565 | 22 | 0.000655 |
|  | **sites** | 1047 | 84 | 0.080229 | 964 | 15 | 0.015560 |
|  |  | **All Regions** |  |  |  |  |  |
|  |  | 228087 | 234 | 0.001026 | 89570 | 54 | 0.000603 |
| **HBCC** |  | **CGI sites** |  |  |  |  |  |
|  | **SV** | 37300 | 14 | 0.000375 | 6643 | 1 | 0.000151 |
|  | **SNV** | 499224 | 23 | 0.000046 | 161836 | 4 | 0.000025 |
|  | **Total** | 536524 | 47 | 0.000088 | 146000 | 5 | 0.000034 |
|  | **sites** | 1586 | 37 | 0.023329 | 860 | 3 | 0.003488 |
|  |  | **Promoters** |  |  |  |  |  |
|  | **SV** | 30605 | 35 | 0.001144 | 2542 | 0 | 0.000000 |
|  | **SNV** | 480520 | 20 | 0.000042 | 56806 | 0 | 0.000000 |
|  | **Total** | 697603 | 71 | 0.000102 | 53426 | 0 | 0.000000 |
|  | **sites** | 1722 | 55 | 0.031940 | 447 | 0 | 0.000000 |
|  |  | **Gene bodies** |  |  |  |  |  |
|  | **SV** | 40669 | 45 | 0.001106 | 11062 | 3 | 0.000271 |
|  | **SNV** | 656934 | 24 | 0.000037 | 209978 | 2 | 0.000010 |
|  | **Total** | 511125 | 99 | 0.000194 | 207949 | 5 | 0.000024 |
|  | **sites** | 2360 | 69 | 0.029237 | 1598 | 5 | 0.003129 |
|  |  | **All Regions** |  |  |  |  |  |
|  |  | 1745252 | 217 | 0.000124 | 407375 | 10 | 0.000025 |

Across the genome, phased methylation data markedly increased effect sizes and enabled the discovery of cis-acting variants with strong regulatory effects on DNA methylation. For example, a 524 bp insertion within a CpG Island on the q-arm of chr 12 is associated with a 20-30% reduction in average methylation of that CGI (Figure 6c). The inserted sequence is likely an expansion of the CGI sequence motif, resulting in fewer reference-coordinate CpGs as they are moved into the insertion and reduced methylation over the reference CGI coordinates in carriers. Notably, this association was not identified by the traditional unphased tensorQTL analysis. In fact, using phased methylation data ASM-LR identified 132 loci where allele-specific methylation effects were masked in unphased tensorQTL models; that is 13% of the 1014 ASM-LR and tensorQTL regions and variants tested by both methods and 28% of the 451 total ASM-LR mQTLs (see methods). 71%, 325, of the ASM-LR mQTLs were also tensorQTL mQTLs, though the hits had different associated variants. On the other hand, out of the 1014 ASM-LR and tensorQTL regions and variants tested by both methods, 557 of those mQTLs were only identified in the unphased tensorQTL model only. Many of the tensorQTL only mQTLs, however, identified extremely small changes in methylation, often less than a 1% change. Due to variable input filtering steps, different variant sets were used for ASM-LR and tensorQTL, which contributes to the lower than expected overlap between the two methods. Again we saw a decrease in the number of significant mQTLs in the ASM-LR runs using deletion filtered methylation data totalling to 64 mQTLs across cohorts and regions. The reduced number of deletion filtered ASM-LR mQTLs is explained in part by the smaller number of tests run on these data, a smaller number of tests that is not confounded by deleted CpGs (Table 3). Similarly there were no HBCC promoter mQTLs that survived multiple test corrections reflecting the reduced number of methylation regions remaining after removing CpG sites affected by deletions (Supplementary Figure 7).

ASM-LR also revealed regions with moderate yet biologically meaningful methylation changes missed by unphased tensorQTL analyses. In the HBCC cohort, we identified a ∼5% increase in methylation across the 1 kb *CR1L* gene body in alleles harboring a deleted CpG island (Figure 6d). Loss of this hypomethylated CGI increases the average methylation level across the gene body. In NABEC, however, a SNV upstream of *CR1L* was the most significant variant associated with a slight increase in methylation across the gene. *CR1L* (complement C3b/C4b receptor 1-like) is involved in cellular responses to hypoxia and cytotoxicity, and altered methylation of this gene has been linked to abnormal cerebral cortex morphology and AD^54,55^. This association was not reproduced in the ASM-LR run with deletions filtered methylation data. ASM-LR was run on 1Mb windows around genes associated with AD and PD in previous literature and we observed that the highest number of ASM-LR mQTLs were in the window surrounding Fibrosin Like 1 (FBRSL1) interestingly 33 of the 40 mQTLs in this window had SVs as their lead variant. Two mQTLs in the FBRSL1 window are near the DEAD-box helicase 51 (DDX51) gene. One being a CpG Island mQTl just upstream of DDX51 and another being the DDX51 gene body mQTL itself which, in fact, was also found to be an HBCC-SV-eQTL associated with the same 90bp deletion ASM-LR identified as being the lead variant for both these mQTLs. The same 90bp deletion was identified in the deletion filtered ASM-LR NABEC CpG Island mQTL but it was associated with a nearby CpG Island instead.

Across the three sets of regions tested, 325 of ASM-LR mQTLs overlapped tensorQTL loci. One notable example is the ZFP57_1 promoter, where a 227 bp deletion upstream of the promoter (Figure 6e) is associated with marked hypomethylation (∼55% methylation in deletion carriers vs. ∼95% in reference samples; Fig. 6c, bottom). *ZFP57* encodes a protein essential for genomic imprinting during embryogenesis ^56,57^. Further studies will be needed to assess the impact of these SVs on ZFP57 protein function. Collectively, these findings highlight the importance of leveraging long-read data to preserve allele-level resolution and uncover epigenetic mechanisms that remain inaccessible using standard, unphased approaches.

## Discussion

LRS has significantly improved our ability to capture complex genetic variation, such as SVs, phase variants of interest, and provide comprehensive methylation data. SVs, in particular, have been shown to influence gene expression in the human brain and are linked to various neurological conditions, but their impact on methylation has been less studied. This highlights the critical role of long-read sequencing in uncovering the full spectrum of genetic contributions to brain function and disease, as it enables the detection of a broader range of genetic variants than traditional genotyping approaches ^59^.

In this study, we present the first brain tissue based comprehensive long-read genomic resource derived from 351 human frontal cortex samples, offering a detailed catalog of 234,905 SVs, 34,087,867 small variants, and DNA methylation profiles across >28 million individual CpG sites. By integrating this dataset with matched short-read RNA sequencing data, we identified numerous SVs as top candidate causal variants influencing both gene expression and DNA methylation in the human brain. These findings provide new insights into the genetic and epigenetic architecture of the brain, demonstrating the power of long-read sequencing to reveal complex regulatory mechanisms. Importantly all data have been made accessible through cloud based analysis platforms to support further research into the functional role of SVs in neurodevelopment and neurodegeneration.

We produced highly contiguous and near-complete locally phased Shasta-Hapdup assemblies for 361 samples covering all but 6% of the GRCh38 reference. Although these assemblies are less contiguous than the multi-technology Human Pangenome Reference Consortium (HPRC) assemblies which have an average phased NG50 of 40Mb, we are able to produce these at scale in a fraction of the time using only one data type^14,60^. Notably compared to ONT sequencing and Shasta Hapdup assembly of 1000 Genomes Project (1KG) cell lines our assemblies of primary tissue from patients are as complete (93% complete; ∼22-35Mbp NG50s to GRCh38) and nearly as contiguous^25^. The assemblies have high base level accuracy, and although highly identical duplicated genes are a challenge for Shasta, we were able to assemble 96% of genes which facilitated the calling of assembly-based SVs ranging from 50bp - 1Mbp. The assembly phase blocks are low, in comparison to HPRC fully phased assemblies, which is a current limitation of phasing without trio information, though as the length and quality of ONT reads increase we expect this to increase.

Our results demonstrate the power of integrating SV data with SNVs, where joint-QTL analysis and fine-mapping identified numerous SVs as likely causal variants influencing gene expression and methylation differences in the brain, variants that would have been overlooked using SRS. By overlapping methylation and expression results, we dissected complex regulatory mechanisms, exemplified by the *NLRP7/NLRP2* locus. At this region, a 1,090 bp deletion of the *NLRP2* promoter, flanked by two *Alu* transposable elements, emerged as a top hit in both mQTL and eQTL analysis. This deletion was correlated with increased *NLRP2* expression in the brain and hypermethylation of a CpG site within exon 6. Interestingly, these findings challenge the conventional view that promoter hypermethylation universally suppresses transcription. In this case, methylation appears to facilitate gene expression, likely through mechanisms involving chromatin structure, allele-specific methylation, or enhancer-promoter interactions. This underscores the context-specific regulatory roles of methylation and highlights the value of integrative analyses for uncovering atypical but biologically significant relationships. To assess the effectiveness of LRS in identifying causal SVs, we compared QTL results from LRS with those from SRS in the NABEC cohort. LRS identified more than twice the number of SV-associated QTLs compared to SRS (740 vs. 358; Supplementary Table 23). For example, the 1,090 bp deletion at the *NLRP7/NLRP2* locus, which LRS accurately resolved, was linked to changes in gene expression and DNA methylation but was undetectable using SRS. Similarly, an insertion associated with *TEX29* expression changes was identified through LRS but missed by SRS. These results demonstrate the superior ability of LRS to resolve complex genomic variations, advancing our understanding of the genetic and epigenetic mechanisms underlying brain function and disease.

Despite these advantages, SV genotyping accuracy remains lower than SNVs due to the technical challenges in merging SV calls across different callers and samples. As a result, we believe the number of SVs identified as causal variants in fine-mapping analyses may be underestimated i.e. if an SV is genotyped less accurately than the small variants included in the testing, or if the SV is inherently variable such that its different suballeles are not merged, it could be excluded even if it is the true causal variant. Additionally, the eQTL analyzes in this study were based on short-read RNA sequencing data, which is often limited to gene-level resolution. Given that SVs can drive differential transcript-level expression and even generate novel transcripts, fully capturing their functional impact will require large-scale long-read RNA datasets which is the natural next step.

To our knowledge, this study represents the first large-scale effort to map methylation QTLs in human brain tissue using LRS and provides the most comprehensive methylation dataset from primary brain to date. It is well-established that genetic variation influences DNA methylation and its relationship with gene expression^61^. We provide the most comprehensive methylation dataset derived from primary brain tissue to date. While the influence of genetic variation on DNA methylation and gene expression is well-established, our work demonstrates the unique advantages of long-read methylation profiling. ONT-based methylation calling has been applied in cancer and neurological disease models ^62,63^, but our study extends this approach to genome-wide mQTL mapping at scale in the human brain.

A key advantage of our study is ASM-LR, a framework for allele-specific methylation QTL analysis from LRS data. Conventional unphased approaches average methylation across alleles, masking haplotype-specific effects. In contrast, ASM-LR preserves allele-level resolution and uncovered 132 loci with allele-specific methylation patterns that were missed by unphased models. Extrapolating that figure across the whole genome we estimate ASM-LR may identify an additional 1,500 (13%) of mQTL hits to the 12,941 mQTLs identified by tensorQTL using unphased data. ASM-LR also revealed increased effect sizes and identified both large-effect cis-regulatory variants (e.g., a 500 bp insertion reducing methylation within a CpG island on chr12) and more subtle allele-specific changes (e.g., deletion-linked methylation shifts in *CR1L* and hypomethylation of the *ZFP57* promoter). These findings demonstrate that haplotype-resolved methylation analysis substantially increases the sensitivity and interpretability of mQTL studies, highlighting the broader value of LRS for dissecting complex epigenetic regulation in the brain.

We were also able to correlate methylation signals with age across samples. It has been shown that changes in DNA methylation have been correlated with age and we were able to recapitulate many of these results which have mostly been identified using targeted arrays and bisulfite sequencing ^64,65^. ONT methylation data is on par with the quality and coverage of whole genome bisulfite sequencing, the historical gold standard for methylation detection, or the targeted approach using whole genome bisulfite sequencing and Illumina BeadChip Arrays ^42,44^. ONT, however, can simultaneously capture genomic variants and measure all modifications on native DNA molecules. We identified many regions that are loosely associated with age and examined those with the largest change over age. The *PCDHG* gene cluster had the most regions that showed increased methylation fractions with age. These genes were not identified as age-related using Infinium HumanMethylation27 BeadChip targeted methylation sequencing, which is an example of how LRS-derived methylation data is more comprehensive for profiling primary tissues^64,66^. Decreases in methylation with age were less common, but similar to prior studies, we did observe a drop in methylation at *FKBP5* ^67^. By developing a methylation catalog of normal brain and prefrontal cortex tissue we are facilitating future work evaluating allele-specific mQTLs and disease-specific differential methylation patterns.

This study has several important limitations, many of which arise from the challenges of applying rapidly evolving technology at a large scale. For instance, after sequencing the NABEC cohort using R9, ONT released new flow cells and chemistries. As a result, the HBCC cohort was sequenced using R10 which offers improved read accuracy but made cross-cohort analyses more difficult. This discrepancy in sequencing platforms may affect the consistency of variant calling between cohorts, highlighting the difficulties of maintaining uniformity in large-scale projects as technology advances. The variability in R9 methylation data compared to R10 has been documented using both chemistries on the benchmark sample HG002 and may be contributing to the low overlap in mQTL hits, especially considering the change in methylation is often quite subtle^44^. Additionally, more sophisticated approaches are needed to improve SV merging methods. This is a fundamentally difficult problem because functionally equivalent SV alleles frequently vary due to independent recurrence and subsequent further acquired variation. We expect that pangenome alignment models will make it easier to group SVs by haplotypic context and features, leading to better SV sets to support downstream association analyses, and the enhancement of allele-specific methylation and expression analyses. Further, while statistical fine-mapping is an effective approach for identifying likely causal variants, functional follow-up studies are necessary to validate their biological impact and confirm their roles in gene regulation and disease mechanisms. Lastly, while this study focused on common germline variation affecting gene expression and DNA methylation in the human brain, future research will explore the role of rare variants, including population-level rare variants and those unique to individual samples, such as low-frequency somatic or mosaic variants.

In summary, we present a foundational long-read genomic and methylation resource derived from 351 neurologically normal brain samples as part of the NIH Center for Alzheimer’s and Related Dementias (CARD) initiative. This dataset represents the first phase of a larger effort to sequence thousands of brains from ancestrally diverse populations, including Alzheimer’s disease and related dementias (ADRD). A key outcome of this work is ASM-LR, which enables robust detection of allele-specific DNA methylation from long-read data. Using ASM-LR, we identified haplotype-specific methylation signatures at key neurodegenerative disease loci, revealing cis-regulatory effects that are undetectable with conventional approaches. Collectively, these data and methods provide a blueprint for future studies aimed at dissecting the interplay between genetic variation, DNA methylation, and gene regulation in the human brain, with the potential to advance mechanistic insights into neurodegenerative disease.

## Methods

### Ethics oversight

The NABEC study was originally approved by the Joint Addiction, Aging, and Mental Health Data Access Committee and more information can be found at the dbGaP website under the study accession ID phs001300.v4.p1. The NIH considered research using post-mortem material as nonhuman subject research and therefore no additional institutional review board approval was required. For the HBCC, brain specimens were obtained under protocols approved by the CNS IRB (NCT03092687), with the permission of the next-of-kin (NOK) through the Offices of the Chief Medical Examiners (MEOs) in the District of Columbia, Northern Virginia and Central Virginia. More information can be found at the NDA website (https://nda.nih.gov/edit_collection.html?id=3151).

### Sample Collection

For the NABEC brain samples, frozen tissues were samples from the frontal cortex for 205 neurologically normal individuals. All NABEC were obtained from three brain banks, the Banner Sun Health Research Institute(https://www.bannerhealth.com/services/research/locations/sun-health-institute/programs/body-donation/tissue), the University of Kentucky Alzheimer’s Disease Center Brain Bank (https://medicine.uky.edu/centers/sbcoa/data-sample-request) and the University of Maryland Brain and Tissue bank (https://www.medschool.umaryland.edu/btbank/). All individuals were of European ancestry and had no clinical history of neurological or cerebrovascular disease, or a diagnosis of cognitive impairment during life. Demographics, tissue source for each subject are shown in Supplementary Table S2. Average age at death was 52.67 yr (range, 15-96 yr), and 36.10% were female.

The HBCC is a brain bank within the National Institute of Mental Health (NIMH) Intramural Research Program. For the HBCC brain samples frozen tissues were dissected from the frontal cortex for 154 neurologically normal individuals. All HBCC samples were obtained from https://neurobiobank.nih.gov/ and data on this can be found at https://nda.nih.gov/edit_collection.html?id=3151 . All individuals were of African or African admixed ancestry and had no clinical history of neurological disease, or a diagnosis of cognitive impairment during life. All specimens examined by a neuropathologist and no degenerative changes were found. Demographics, tissue source and for each subject are shown in Supplementary Table S2. Average age at death was 44.22 yr (range, 18-85 yr), and 36.36 % were female.

### DNA processing

The DNA processing ^68,69,70^ protocols are explained in detail and are publicly available on the protocols.io platform. In brief, ∼40mg of frozen frontal cortex tissue was homogenized with a Tissueruptor instrument (Qiagen). High molecular weight (HMW) DNA was then extracted using a Kingfisher Apex instrument (Thermofisher) with a custom script and Nanobind Tissue Big DNA kit, which uses 3mm Nanobind disks (Circulomics/PacBio, US). The HMW DNA was sheared to a target size of 30kb with the DNA Fluid + needles at speed 45 for two cycles on a Megaruptor3 instrument (Diagenode).

### PromethION Long-read sequencing

For the NABEC cohort libraries were constructed using an SQK-LSK 110 kit (ONT). To ensure minimal DNA loss during library preparation, and to retain long DNA fragments, the following modifications were made to the standard SQK-LSK 110 (ONT) protocol; 1) 4.5 ug of DNA was used as starting input, 2) to reach the 4.5ug DNA input the starting DNA volume was usually higher than the recommended 47ul. In this case, the volume of AMPure XP beads was modified to match the input DNA volume, i.e. if 58ul of DNA was added then 70ul was added of AMPure XP beads, 3) during DNA repair and end-prep, 75% ethanol was used for washing rather than 70%, 4) at step 16 of DNA repair and end-prep, the original elution time was 2 minutes at room temperature. This was modified to 3 minutes at 37°C with light shaking on a Thermomixer instrument (Eppendorf) at 450rpm, 5) during Adapter ligation and clean-up, 45uL of AMPure XP beads were used 6) SFB was used, and finally 7) in step 16 of the Adapter ligation and clean-up, the final elution conditions were changed to 20 minutes at 37°C. For HBCC libraries were constructed using an SQK-LSK 114 kit (ONT) using the same modifications as above however where 2.5 ug of DNA was used as starting input.

PromethION sequencing was performed as per manufacturer’s guidelines (ONT, FLO-PRO002) with minor adjustments, such as for R.9 400ng of the library was loaded onto each primed R9.4.1 flow cell to maximize data output and for R.10 180ng of the library was loaded onto each primed R10.4.1 flow cell. Each sample required at least two or three loads to achieve > 115GB total data output over 72 hours.

### Basecalling and Alignment

All samples were basecalled and aligned with minimap2 on the NIH HPC Biowulf cluster. NABEC Basecalling was performed using Guppy v 6.12 and HBCC basecalling was performed using Guppy v 6.38.

### Small variant calling within NAPU

As a part of the NAPU workflow small variants (substitutions and indels smaller than 50bps) are called from ONT reads using either P.E.P.P.E.R-Margin-DeepVariant^71^ or DeepVariant^72,73^ (ONT_R104 model) for the R9 and R10 cohorts respectively. Small variant gVCF files, produced by DV, were then merged within cohorts using glnexus and the “DeepVariantWGS” configuration, for R9 “NoCall” gVCF entries were removed prior to merging^74^. Illumina short-read whole genome sequencing data for NABEC was obtained from dbGaP (Accession phs002636.v3.p1).

### Structural variant calling

All brain samples were processed with the NAPU workflow which uses hapdiff and Sniffles2 to produce assembly and read based structural variant calls respectively ^14,15^. Read based, reference-free, phased SVs were called using Sniffles2 v.2.3^15^. For assembly based structural variants diploid de novo assemblies are created by first making a haploid Shasta assembly and then using Hapdup v.0.12 to locally phase contigs to create a diploid assembly^14,20^. The diploid assemblies are then aligned with minimap2 (v 2.23-r1111)^75^ to the GRCh38 reference and SVs are called with Hapdiff ^14^. We used Shasta v0.11.1^20,76^ with the “Nanopore-CARD-Jan2022.conf” configuration file for R9 samples and “Nanopore-R10-Fast-Nov2022.conf” configuration with these command line arguments "--Reads.minReadLength 8000 --Align.minAlignedMarkerCount 500 --Align.minAlignedFraction 0.75" to account for the shorter read lengths observed in R10 sequenced brain samples. For samples with high read coverage we also used the “--Reads.desiredCoverage 150Gbp” which removes the shortest reads until the coverage is below the desired coverage limit. Samples sequenced with R10 had lower coverage, read lengths, and identity relative to the R9 samples, so to produce more contiguous Shasta assemblies, we relaxed the R10 length and alignment criteria (Supplementary Figure 1A, 1B). We attribute the reduced quality in R10 sequence brain samples to be due to reduced quality brain tissue samples instead of due to the R10 chemistry.

SVs were merged across all samples within a cohort as well as across cohorts using either Sniffles2 or Truvari^77^. Sniffles2 was used to merge and genotype read-based SVs called by Sniffles. Truvari merge with a 75% similarity and size cutoff was used to merge hapdiff SVs within cohorts. To merge hapdiff variants, first bcftools^78^ is used to combine all vcf entries for all cohort samples which was 1.17 million NABEC SVs and 1.07 million HBCC SVs. Truvari then merged similar SVs within those sample combined VCFs which collapsed the number of NABEC SVs to 165,854 and HBCC to 210,808 SVs. After merging SVs within a cohort we used assembly alignment coverage beds created by hapdiff to then assign reference genotypes (0/0) to assembly-based SVs where there was assembly coverage but no SV called. Truvari is not currently capable of joint genotyping, so in order to include hapdiff SVs in eQTL SV analysis, which filters by genotype, we performed this manual reference genotype assignment. Sniffles2 v2.3, on the other hand, performs its own merging and joint genotyping, which resulted in 213,279 NABEC and 210,163 HBCC SVs. Sniffles2 v2.3 did take substantially longer to merge and joint genotype at over 7 and 5 hours for each cohort. Sniffles, however, is only able to merge calls it produces itself (requiring a snf file it produces during variant calling) thus, Truvari was used to merge variants between SV callers and then the two cohorts.

In order for Truvari to merge SVs from Hapdiff and Sniffles within each cohort we had to do some curation of the Hapdiff+Truvari merged cohort VCF and the Sniffles merged VCF. Truvari compares sequence similarity so the symbolic representation of Sniffles SVs was replaced with reference sequence information using a script provided by Truvari (https://github.com/ACEnglish/truvari/discussions/216). Hapdiff was able to capture the individual variation of each sample but after merging this resulted in many alleles with a small number of sample genotypes per variant. Variants with a minor allele frequency less than 5% were filtered out prior to running the QTL. For this reason we resorted to choosing Sniffles variants by setting Hapdiff quality scores to 0, since it is unable to assign a quality score to its’ assembly based SV calls, and then kept the SV representation with the highest quality score. Variants from each SV caller were combined using bcftools concat, summing to 366,717 NABEC SVs and 436,838 HBCC SVs, and after Truvari merging at 75% similarity there were 193,601 and 241,202 NABEC and HBCC cohort SVs respectively.

To ensure consistency of SV identification across both cohorts and enable comparison of allele frequencies and QTL results, we merged variants from both cohorts to create a unified set of variants. First bcftools combined samples from both cohorts and then Truvari collapse was used again to combine similar variants at 75% similarity and a max size of 60Kbp. After this merge there were 278,732 variants but over 58 thousand were still contained multi-allelic variants. These multi-allelic variants were extracted and further merged to combine larger variants over larger reference distances as well as combine insertions and deletions at the same loci. Merging these multi-allelic variants more loosely reduced those 58 thousand multi-allelic variants to just 14,717 representative SVs. These variant sets were once again combined for a final set of 234,906 single allelic variants across both cohorts and variant callers. That is 139,841 NABEC variants and 180,430 HBCC variants. The SVs were compiled into a UCSC Genome Browser track allowing visualization alongside other genomic features like promoter and CpG Island tracks. The track is available at (https://genome.ucsc.edu/cgi-bin/hgTrackUi?g=cardSv).

### Methylation analysis

Phased methylation beds for both R9 and R10 cohorts were created using Modkit pileup. We ran Modkit using the GRCh38 reference, restricted to 5-methylcytosine (CpG) motifs and combined across read strands. We then created aggregated regional methylation bed files using bedtools map and limiting to regions with minimum 10 CpG sites and 5 reads (https://github.com/nanoporegenomics/napu_wf/blob/r10/wdl/workflows/bedtoolsMap.wdl). We included this raw methylation data as well as deletion filtered methylation data for mQTL analysis. To ensure that we use an equal number of CpGs in all samples, CpGs within deletions were removed from all samples. To do this deletions were isolated from a merged set of cohort SVs, converted into a bed file, and used to remove CpGs within cohort deletions from the cohort methylation data prior to aggregating. We averaged methylation over CpG sites across gene promoter regions, gene bodies, and CpG Islands. We obtained the gene promoters region bed track from the Eukaryotic Promoter Database^79^. The promoter regions in the database are 60 bps for this work we extended them up and down stream by 1kb. Gene body regions were from the UCSC genome annotation database for the Dec. 2013 (GRCh38/hg38) assembly of the human genome (hg38, GRCh38 Genome Reference Consortium Human Reference 38 (GCA_000001405.15); https://hgdownload.cse.ucsc.edu/goldenpath/hg38/database/refFlat.txt.gz). To prevent multiple entries for genes with multiple isoforms the isoform with the most exons was chosen as the representative length of the gene body. CGI bedtrack is from the UCSC genome browser CpG Island track.

We used the available cohort meta-data to examine regional methylation which correlates with age in the brain. To do this we ran linear regressions on autosomal CGI’s, and whole genome windows (1kb, 50 CpG min) with post-mortem interval and sex as covariates over the sample age. After Benjamini-Hochberg multiple test correction, 28%, 145,086, whole genome windows were found to be significantly associated with age (FDR < 0.05). Of the UCSC Genome Browser Track autosomal CpG Islands 14%, 3,890, and 23%, 5,778, in NABEC and HBCC respectively were found to be significantly associated with age (FDR < 0.05).

Next we aimed to correlate methylation levels with genetic variants and performed mQTL analysis. For use in the mQTL we averaged methylation at all sites in regions of interest: promoters, gene bodies, and CpG Islands. These aggregated methylation regions were used as the input for mQTL analysis. Of the 29,598 promoters 94% NABEC and 93.5% HBCC promoters remaining after filtering for read coverage and CpG density with 27,465 (92.7%) confidently covered by both cohorts. After filtering, the NABEC and HBCC cohorts retained 93.4% and 91.4% of the 27,949 CGIs in the UCSC Genome Browser Track for a total of 90.1% of CGIs covered by both. Gene body regions were defined as 62,700 unique gene entries from Gencode V.44. After filtering these regions for coverage and minimum number of CpGs 65.3% of gene bodies were covered by both cohorts. Individually the NABEC cohort retained 67% of these gene bodies and HBCC kept 66%.

### Structural variant annotation

STIX [10.1101/2024.09.30.615931 and 10.1038/s41592-022-01423-4] was used to annotate the harmonized SV callset. STIX annotates all SVs with a long-read index, which includes 1,108 samples (1,008 samples from the 1000 Genomes Project[10.1093/nar/gkz836] and 100 samples from the HPRC project[10.1038/s41586-023-05896-x]) across 5 super-populations. We used ‘-s 100’ and ‘-T 5’ to set a padding base of 100 and required a minimum of 5 reads to define true signals. STIX long-read indices were built using excord-lr to extract raw SV signals from BAM or CRAM files. Giggle [10.1038/nmeth.4556] was used alongside STIX to create indices. All related tools for index creation and annotation can be found in the repository (https://github.com/zhengxinchang/stix-suite) with version 1.0.1. A total of 234,906 variants were annotated, of which 9,747 failed annotation because STIX does not index SV signals in random contigs. As a result, 225,159 variants were annotated successfully.

### Ancestry analysis

Ancestry predictions for 11 different ancestry groups were rendered using GenoTools^18^ ancestry pipeline with a serialized model trained on variants in common between the NeuroBooster Array^80^ and the reference panel outlined in GenoTools. This method utilizes PCA and UMAP transformations and an XGBoost classifier which targets several superpopulations and selects subpopulations relevant to Neurodegenerative diseases.

### Population level allele frequency comparison

Genotype counts from the NABEC+HBCC, NABEC, and HBCC callsets were used to calculate variant frequencies, which were compared with STIX frequency annotations^31^. Homozygous and heterozygous genotypes were collectively categorized as "mutated," while reference genotypes (0/0) were assigned a frequency of zero. The variant frequency was defined as the proportion of mutated samples (homozygous or heterozygous) among the total number of samples with valid genotypes (excluding missing genotypes). To ensure robust frequency estimates, minimum genotype thresholds were applied: 350 samples for NABEC+HBCC, 200 samples for NABEC, and 140 samples for HBCC. Pearson correlation coefficients were calculated to evaluate the concordance between STIX annotations and variant frequencies observed in the respective callsets.

### Expression analysis

#### NABEC Bulk short-read RNA sequencing data

To identify the functional impact of the new genetic variants, sample data was taken from the North American Brain Expression Consortium (NABEC; <u>dbGaP Accession phs001300.v4.p1</u>). Bulk RNA-seq data from all samples was quantified using Salmon ^81^ with the filtered Gencode v43 human transcriptome index. Bulk frontal cortex expression from NABEC contained 206 samples covering 60880 quantified genes. Sex specific genes were subsetted and plotted to identify any sample sex mismatches, and for brain samples. 34102 genes with a missingness over 0.25 were excluded, leaving 26778 well detected genes. To standardize, expression data was quantile transformed and were scaled to 0 to 1.

#### HBCC Bulk short-read RNA sequencing data

To identify the functional impact of the new genetic variants, sample data was taken from the HBCC, https://nda.nih.gov/edit_collection.html?id=3151). Bulk RNA-seq data from all samples was quantified using Salmon ^81^ with the filtered Gencode v32 human transcriptome index. Bulk frontal cortex expression from HBCC contained 86 samples covering 60880 quantified genes. Sex specific genes were subsetted and plotted to identify any sample sex mismatches. 34,201 genes with a missingness over 0.25 were excluded, leaving 26679 well detected genes.

#### Quantitative trait loci analysis

The expression and methylation data was normalized by quantile-normalization and min-max scaled and was used for cis quantitative trait loci (QTL) analysis using tensorQTL.^82^ Sex, age, genetic principal components (PCs), trait PCs, postmortem interval, and group in the datasets (’SH’, ’KEN’, ’UMARY for NABEC) were used as covariates. QTL analyses were analyzed using three variant sets including SVs, SNVs (illumina data in NABEC, DeepVariant SNVs in HBCC), and harmonized SV and SNVs. Genetic PCs were calculated for each variant type and caller using variants within the cis windows around each region of interest. Variants were filtered based on the following criteria: 1) minor allele frequency < 0.05, 2) genotyping rate < 0.95, 3) Hardy-Weinberg equilibrium pvalue > 0.001, 4) non autosomal variants. Genetic PCs were generated in PLINK (v1.9). PCs of gene expression data were calculated using the PCA module from the sklearn.decomposition package. The number of genetic PCs and expression PCs to include in the QTL analysis was selected by determining the point of maximum curvature using kneed (https://github.com/arvkevi/kneed). All variant–phenotype pairs within a specified window (± 1 Mb) around the phenotype were included in the QTL. Significant hits were identified as having a false discovery rate corrected p-value < 0.05 (FDR < 0.05).

#### Fine-mapping of causal variants QTL

We applied CAVIAR to assess LD and identify the most likely causal variant for each eQTL and mQTL. CAVIAR integrates summary statistics with LD information across the associated locus to estimate the causal probability of each variant. QTL were identified with q-value < 0.05. Z-scores were computed for each gene-variant pair using linear regression slope and standard deviation as an input for CAVIAR. For each significant loci, we used Z-scores and LD for the top 100 variants including the best SV (if no SV was included in the top 100 variants) with causal set size of 1 to run CAVIAR. Posterior probabilities were used to evaluate the causality of each variant.^83^

For each of the 2,064 eQTLs in NABEC and 1,142 eQTLs in HBCC, we selected the top 100 variants (ranked by p-value) within a 1 Mb cis window that were most significantly associated with the expression of that gene. CAVIAR was then used to assign a causal likelihood and relative ranking to each of these 100 variants based on the strength and direction of their associations, as well as the pairwise LD structure across the region^38^. Most gene loci analyzed contained both an SV and small variants for CAVIAR to analyze; however, for loci where no SV was present within the 1 Mb window only small variants were evaluated which occurred in 0.5% of NABEC and 1.4% of HBCC eGenes.

#### Short-read SV calling for QTL comparison

To assess whether the LRS QTL hits identified in this study could also be detected using SRS methods, we analyzed existing SRS-derived SV calls for the NABEC cohort, as previously described by Billingsley et al^84^. In brief, SV data were generated from WGS processed through the AMP-PD pipeline. DNA samples were sequenced by either Macrogen or the Uniformed Services University of the Health Sciences (USUHS) using Illumina TruSeq DNA library preparation protocols, resulting in average insert sizes of ∼300–410 bp. Sequencing was conducted on the Illumina HiSeq X platform, and reads were aligned to the GRCh38DH reference genome using the Broad Institute’s standardized functional equivalence pipeline. SVs were identified using the GATK-SV pipeline (https://github.com/broadinstitute/gatk-sv), which integrates multiple SV callers, depth-based analyses, and downstream filtering to ensure high-quality variant discovery. In total 353,426 SRS-derived SV calls were available for 189 NABEC samples.

#### Allele-specific methylation QTL analysis with ASM-LR

To identify genetic variants associated with allele-specific DNA methylation, we developed ASM-LR, a linear modeling framework designed to detect methylation QTLs using phased long-read whole-genome sequencing data. ASM-LR integrates phased SNVs and SVs with methylation information resolved at the haplotype level, enabling the identification of cis-acting genetic effects that are masked by traditional, unphased approaches.

#### ASM-LR Input data and preprocessing

ASM-LR requires four primary inputs. First, it uses phased genetic variants, including both SNVs and SVs, generated using the NAPU pipeline. The NAPU pipeline used margin to coordinate, or harmonize, the genotypes of SNVs and SVs, so that even though variants are called from the reads and/or assembly using different tools the variant genotypes are coordinated to be on the same allele ^14,71^. Second, it incorporates aggregated methylation information, specifically haplotype-resolved methylation levels aggregated across the defined methylation regions in this study; CpG islands (CGIs), promoters, and gene bodies. The phased methylation data is also harmonized to coordinate with the variant genotypes, so a variant and the respective methylation values of that haplotype are reported to be on the same allele. Phased methylation data was merged across haplotypes into a single input table with average methylation per region, sample, and haplotype. Third, the analysis is limited to a curated list of 171 genes implicated in AD or PD, serving as the primary regions of interest. Finally, sample-level covariates, including age, sex, biobank of origin, and the first 20 genetic principal components generated from CpG island, gene body, or promoter-specific SNVs with plink, are included in the model to account for confounding factors. Data input preprocessing for the QTL is further documented in our data standardization GitHub repository (https://github.com/NIH-CARD/CARDlongread_data_standardization).

#### Variant filtering and pruning

Prior to association testing, genetic variants were filtered using PLINK v1.9, applying standard quality control thresholds to ensure reliable association results. Variants were retained if they had a minor allele frequency (MAF) of at least 5%, conformed to Hardy-Weinberg equilibrium (HWE) with p > 1 × 10⁻⁶, and had a genotyping call rate of at least 95%. To reduce redundancy from correlated variants, we performed linkage disequilibrium (LD) pruning using an r² threshold of 0.3 given a 1000 variant window and 50 variant step size (plink v2.00a6LM). Only variants passing these filters and within 500 kb up or downstream of the defined gene set were retained for testing.

#### Association testing

We performed association testing using a linear regression model to evaluate the relationship between genotype and haplotype-resolved methylation levels. For each variant–region pair, the average methylation frequency across the region (CpG island, promoter, or gene body) was modeled as a function of genotype and covariates:

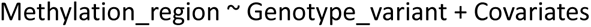

Covariates included age, sex, biobank of origin, and 20 genetic principal components to account for population structure. Association testing was conducted separately within each cohort. During association testing, methylation regions and variants were only considered if the minor allele frequency (MAF) for at least one haplotype was greater than 0.05 if the other haplotype was not present in the methylation data.

#### Significance threshold and prioritization

Variants were considered significantly associated with allele-specific methylation if they passed a Benjamini-Hochberg (B-H) false discovery rate (FDR) correction threshold of <0.05. Loci were then prioritized for follow-up if they overlapped known regulatory elements or fell within regions previously associated with AD/RD or PD through genome-wide association studies (GWAS). The ASM-LR pipeline is the first publicly available tool for genome-wide allele-specific methylation analysis using long-read sequencing data. It is freely accessible on GitHub and designed to support the research community in uncovering haplotype-resolved epigenetic regulation across diverse tissues and populations.

#### Code availability

The complete pipeline implemented in WDL is available at https://github.com/nanoporegenomics/napu_wf. The code used to process and analyze the data for this study is publicly available at https://github.com/NIH-CARD/CARDlongread_NABEC_HBCC_QTL_manuscript, which includes scripts and workflows for structural variant discovery, eQTL and mQTL analyses, and downstream data integration. The ASM-LR pipeline, a standalone tool for allele-specific methylation analysis from long-read data, is available at (https://github.com/molleraj/CARDlongread_allele_specific_QTL) as is the data standardization pipeline to prepare phenotype and genotype data for ASM-LR (https://github.com/NIH-CARD/CARDlongread_data_standardization). Additional details about the computational tools and parameters used in this study are described in the Methods section of the manuscript.

## Data availability

Human brain sequencing datasets are under controlled access and require a dbGap application (phs001300.v4; substudy phs003181.v2.p1) (phs000979.v4,). The data is available through the restricted AnVIL workspace at NABEC: https://explore.anvilproject.org/datasets/a00842c8-12f8-4f92-a7b3-01bb73ffd4e4 and HBCC: https://explore.anvilproject.org/datasets/a4e936d1-d81a-475d-95be-b5cd41de921d, as well as the Alzheimer’s Disease Workbench (ADWB, https://www.alzheimersdata.org/ad-workbench). The SVs UCSC Genome Browser track is available at https://genome.ucsc.edu/cgi-bin/hgTrackUi?g=cardSv.

## Acknowledgements

We thank members of the North American Brain Expression Consortium (NABEC) for providing samples derived from brain tissue. We are grateful to the Banner Sun Health Research Institute Brain and Body Donation Program of Sun City, Arizona for the provision of human biological materials. This work was supported in part by the Intramural Research Program (IRP) of the National Cancer Institute (NCI), the National Human Genome Research Institute (NHGRI, ZIAHG200398), National Institute on Aging (NIA, ZIAAG000534, ZIAAG000538), and the Center for Alzheimer’s and Related Dementias (CARD), within the Intramural Research Program of the NIA and the National Institute of Neurological Disorders and Stroke (NINDS). The HBCC is funded by the National Institute of Mental Health (NIMH)-IRP (ZIC MH002903). DEM is supported by NIH grant DP5OD033357. B.P. was partly supported by NIH grants: R01HG010485, U24HG010262, U24HG011853, OT3HL142481, U01HG010961, and OT2OD033761. M.M. was supported by NIH grant T32HG012344. We acknowledge the support of Oxford Nanopore Technologies staff in generating this data set, in particular A. Markham and J. Anderson. We also acknowledge the support of the PacBio team in generating the wet-lab protocol, in particular K. Liu, J. Burke, M. Kim & D. Kilburn.

This work utilized the computational resources of the NIH STRIDES Initiative (https://cloud.nih.gov) through the Other Transaction agreement - Azure: OT2OD032100, Google Cloud Platform: OT2OD027060, Amazon Web Services: OT2OD027852. This work utilized the computational resources of the NIH HPC Biowulf cluster (https://hpc.nih.gov). Some authors’ participation in this project was part of a competitive contract awarded to DataTecnica LLC by the NIH to support open science research. M.A.N. also owns stock in Character Bio Inc. and Neuron23 Inc. This research was supported in part by the IRP of the NIH. The contributions of the NIH authors are considered Works of the United States Government. The findings and conclusions presented in this paper are those of the authors and do not necessarily reflect the views of the NIH or the U.S. Department of Health and Human Services.

---

**Supplementary Figure 1.**
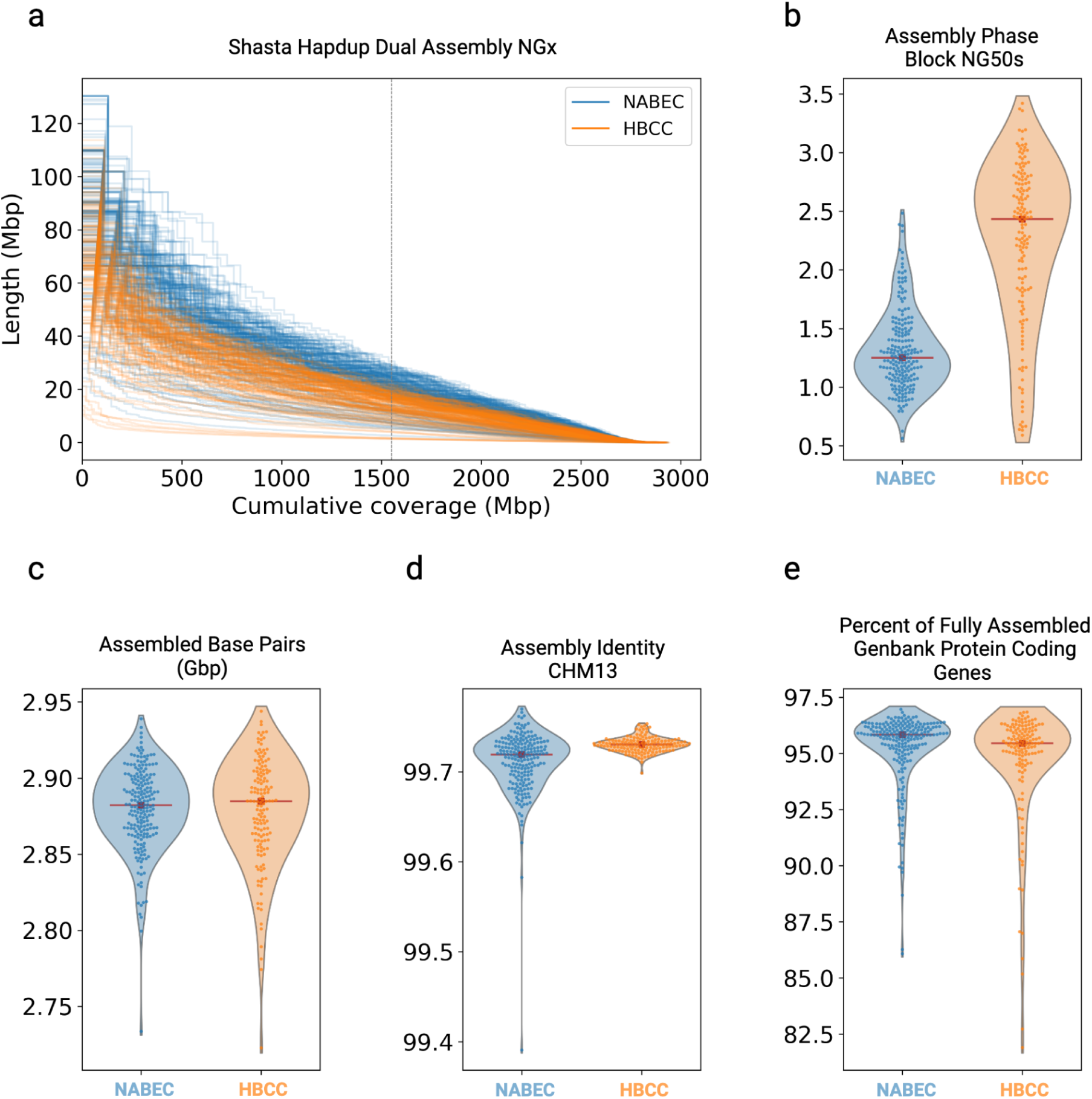
: Assembly statistics for both cohorts. **a.** Shasta+Hapdup Dual assembly NGx plot for HBCC and NABEC Cohorts, compared to the 3.1 Gb length of the T2T-CHM13v2.0 assembly. **b.** NG50’s of phased assemblies illustrate the increased phase block size of African American ancestry HBCC R10 samples. The red line is the median value. **c.** The number of assembled base pairs in dual assemblies aligned to CHM13v2.0. **d.** Contig identity mapped to CHM13. **e.** Percentages of Genbank v44 protein-coding genes completely assembled in each samples’ dual assembly.

**Supplementary Figure 2.**
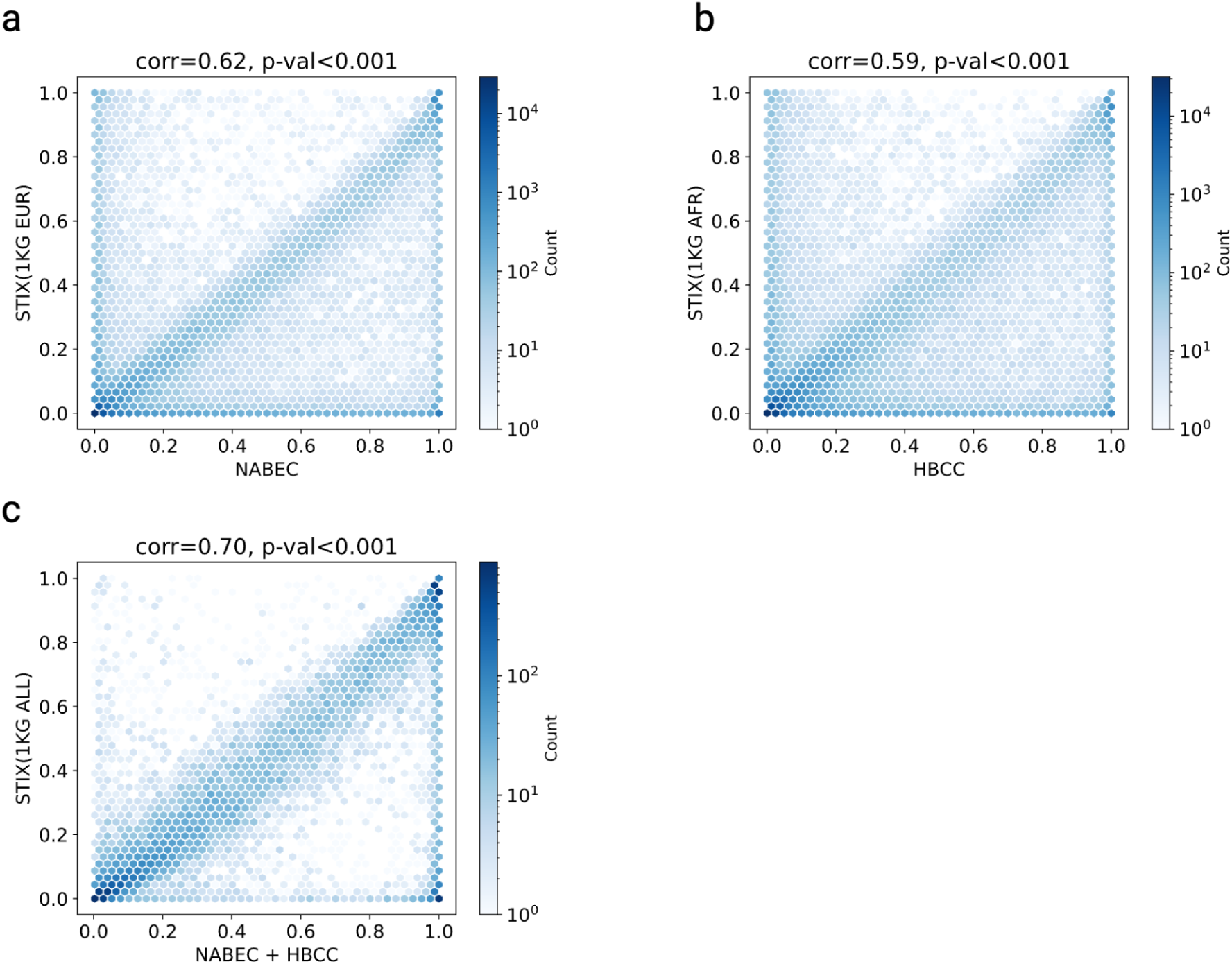
Comparison of STIX frequency and variant frequency within three callsets. Variants with valid genotypes were included. a) NABEC only, minimal genotypes = 200. b) HBCC only, minimal genotypes = 140. c) NABEC + HBCC, minimal genotypes = 350. Pearson correlation was used to evaluate the concordance between the STIX frequency and variant frequency within the callsets. The color indicates the number of variants located in specific hexagonal bins.

**Supplementary Figure 3.**
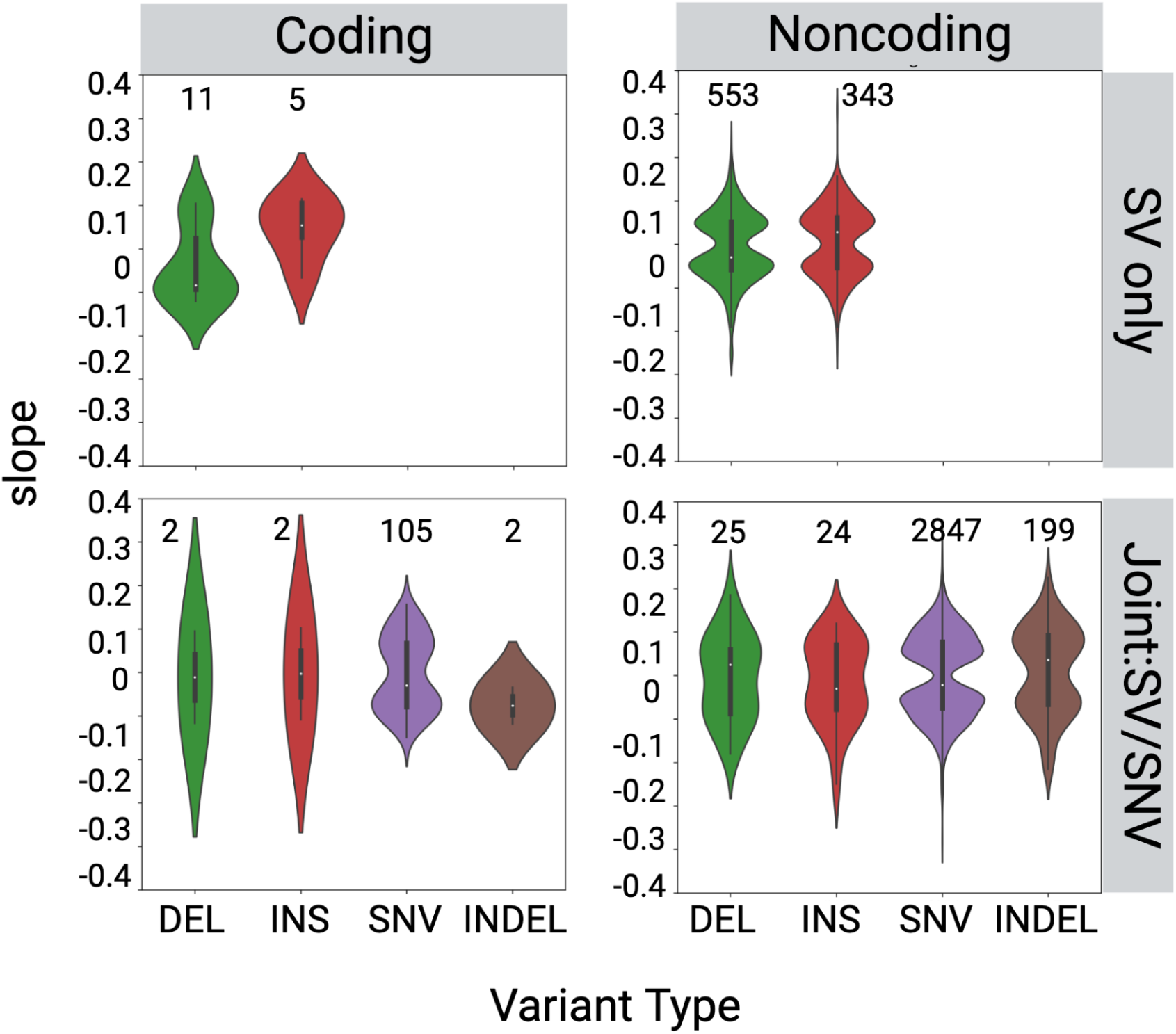
Comparison of effect sizes between variant types for eQTL, left panel. In each panel, panels is divided in four sections, top left SV-eQTL hits in overlapping with coding regions, top right SV-eQTL hits in non-coding regions, bottom left SV-SNV joint eQTL hits in coding regions, bottom right SV-SNV joint eQTL hits in non-coding regions.

**Supplementary Figure 4.**
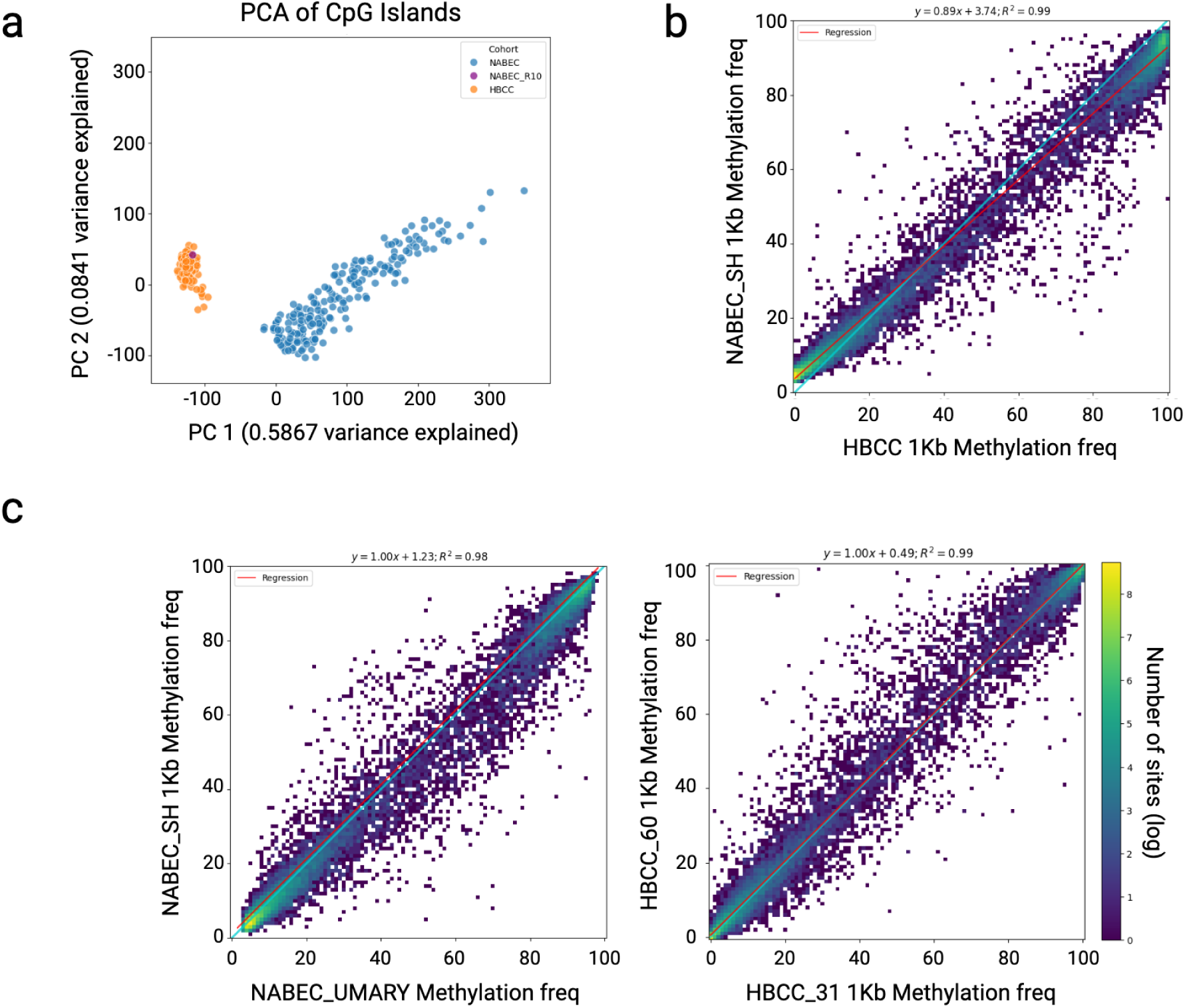
a. PCA of NABEC and HBCC methylation averaged over CpG Islands. One NABEC sample was sequenced with both R9, plotted in blue, and R10, purple. The clustering of the R10 sequenced European NABEC sample with the other R10 sequenced African or African admixed ancestry HBCC samples suggests that the technical differences in methylation frequency are stronger than the ancestry differences. b,c. Heatmaps of pairwise comparisons of autosomal methylation frequency between a NABEC and HBCC sample (left) two NABEC samples, and two HBCC samples. The light green line is plotted on the diagonal and the regression line fitted to the data is plotted in red. In the far left panel, the red line is off the diagonal with respect to the NABEC cohort (y-axis); the regression line equation and the R^2^ correlation are the titles of these panels.

**Supplementary Figure 5.**
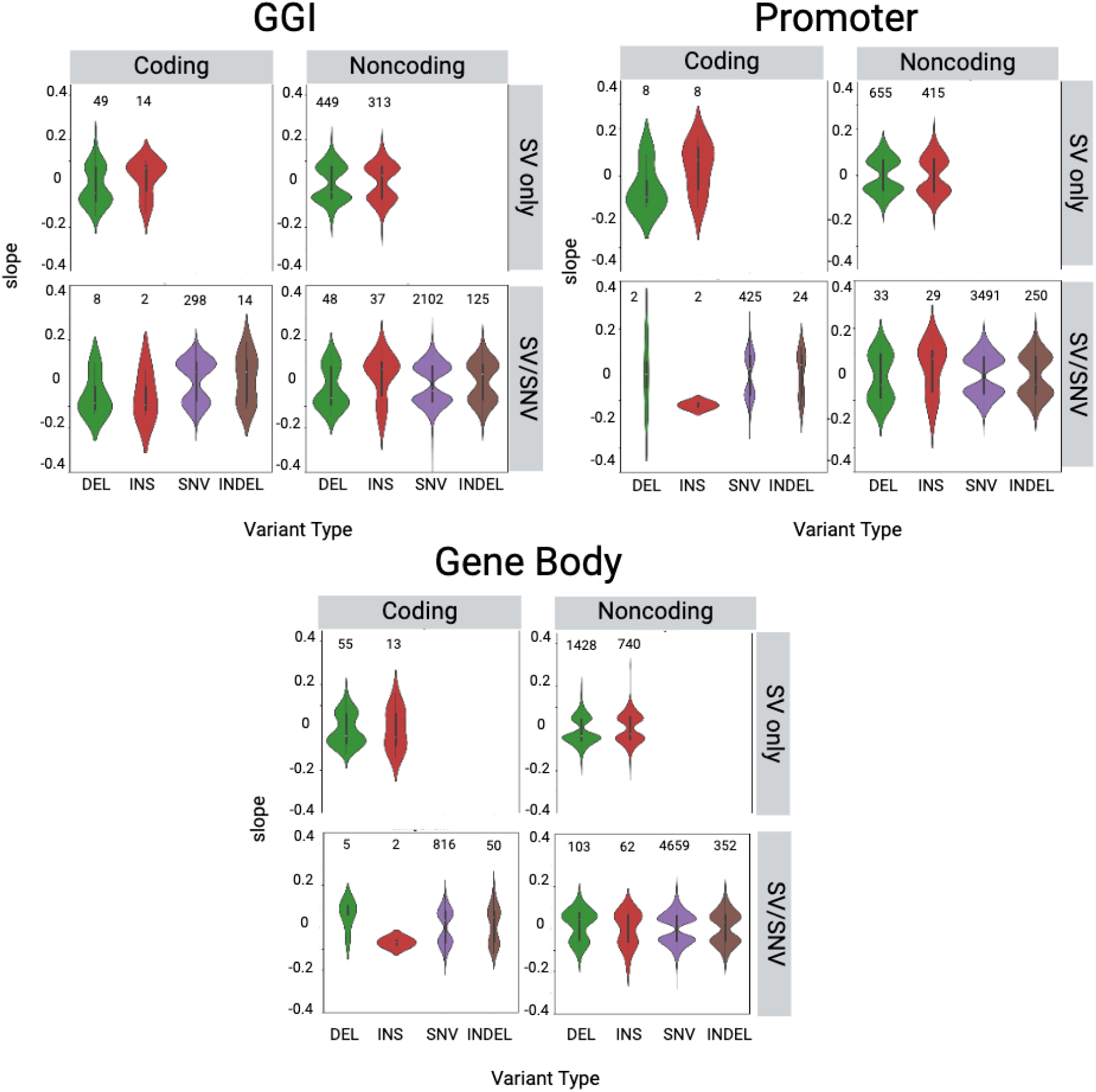
Comparison of effect sizes between variant types for mQTL, left panel; Cpg islands, center panel; promoters, right panel; gene bodies. In each panel, panels is divided in four sections, top left SV-mQTL hits in overlapping with coding regions, top right SV-mQTL hits in non-coding regions, bottom left SV-SNV joint mQTL hits in coding regions, bottom right SV-SNV joint mQTL hits in non-coding regions.

**Supplementary Figure 6.**
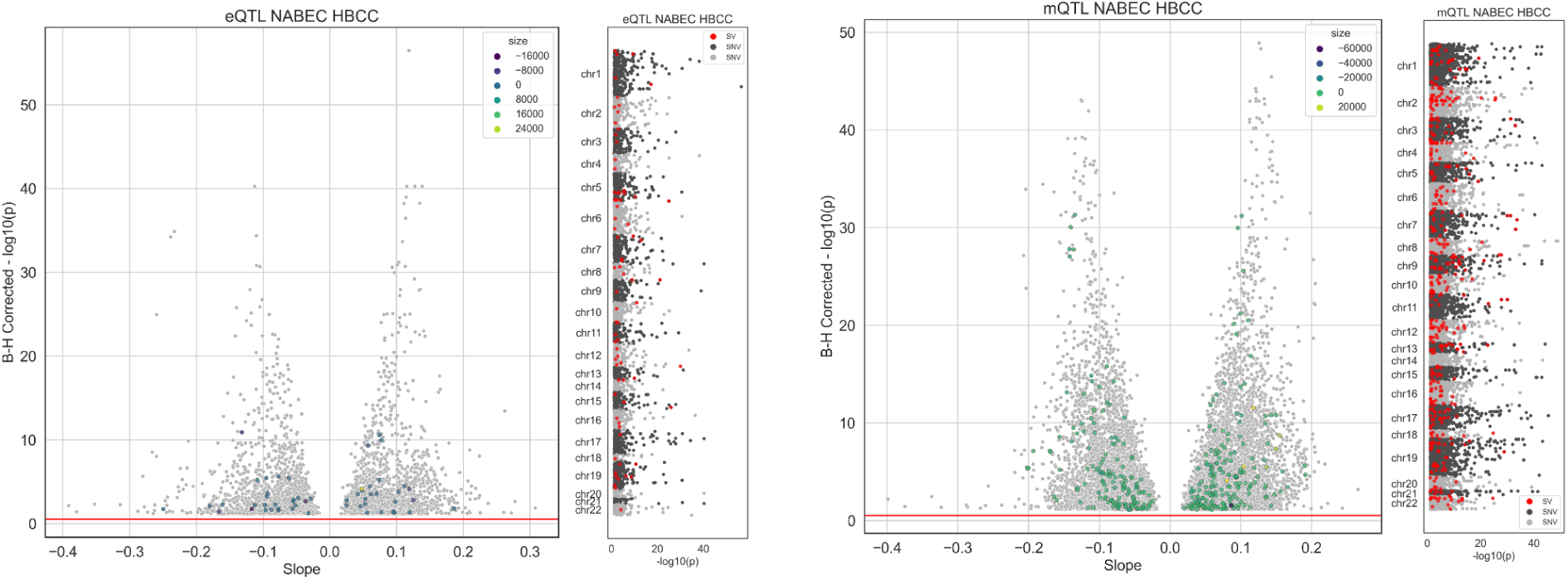
Volcano and Manhattan plots of eQTL and mQTL results combined across all tests for both cohorts. Grey points in both plots represent QTLs with an SNV as the lead variant and colored points represent QTLs with a SV as the top hit. The color of SV led QTLs in the volcano plot are colored by SV size (negative size denotes deletions).

**Supplementary Figure 7.**
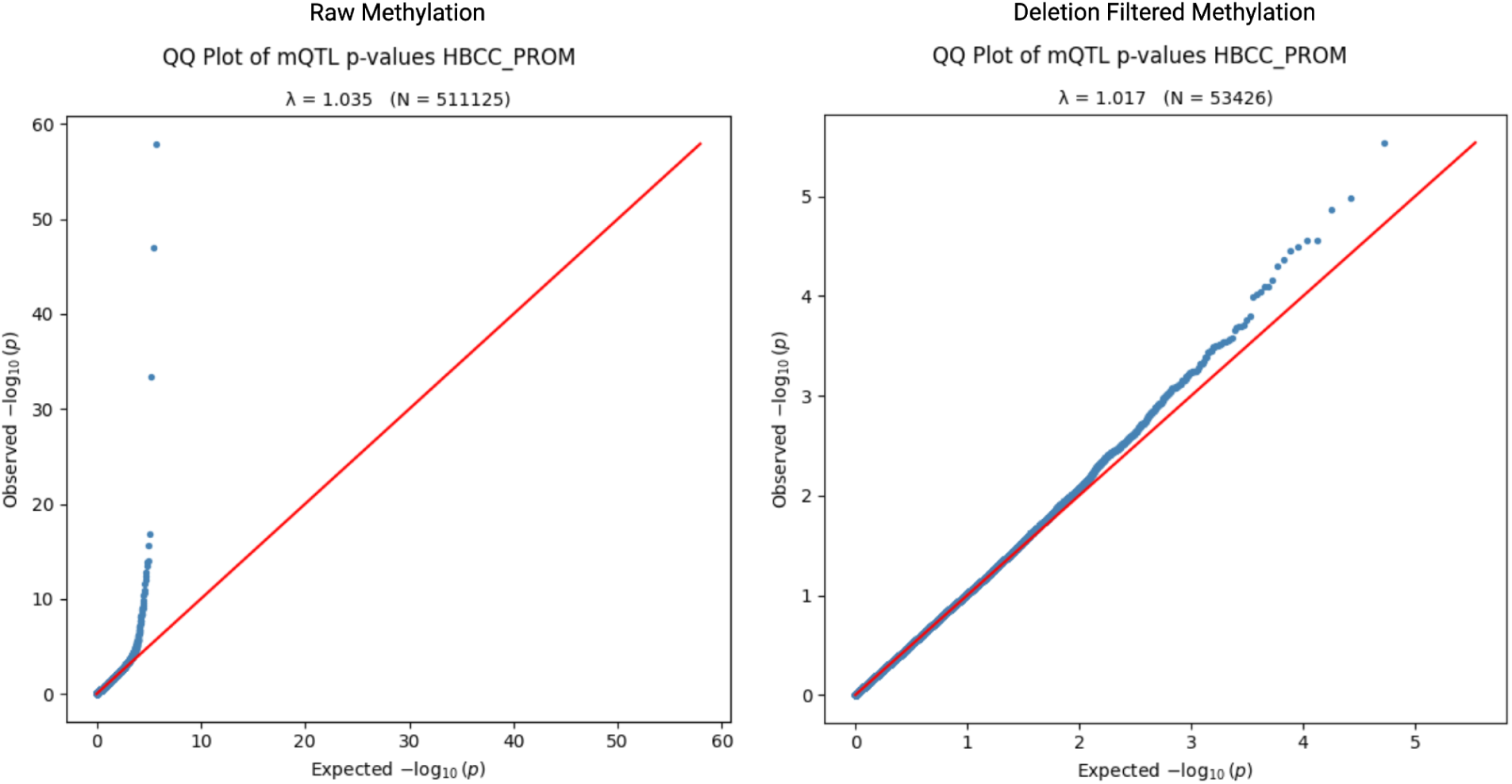
Q-Q Plots of observed p-values versus the expected p-values for the ASM-LR HBCC promoter mQTL. The lefthand plot is for the HBCC promoter mQTL that used unfiltered methylation data and the right plot is for the deletion filtered methylation mQTL analysis, in which CpGs deleted in any sample were excluded prior to promoter-level aggregation. The genomic control (λ) values and the number of tests are in the subtitles of the plot. Although there were no HBCC promoters in the deletion-filtered mQTL that passed multiple test corrections, largely due to the reduced number of tests shown in the deletion filtered mQTL.

## References

1. Ding, W. et al. Adaptive functions of structural variants in human brain development. Sci Adv 10, eadl4600 (2024).

2. Han, L. et al. Functional annotation of rare structural variation in the human brain. Nat Commun 11, 2990 (2020).

3. van Bree, E. J. et al. A hidden layer of structural variation in transposable elements reveals potential genetic modifiers in human disease-risk loci. Genome Res 32, 656–670 (2022).

4. Sleegers, K. et al. APP duplication is sufficient to cause early onset Alzheimer’s dementia with cerebral amyloid angiopathy. Brain 129, 2977–2983 (2006).

5. Singleton, A. B. et al. alpha-Synuclein locus triplication causes Parkinson’s disease. Science 302, 841 (2003).

6. Makino, S. et al. Reduced neuron-specific expression of the TAF1 gene is associated with X-linked dystonia-parkinsonism. Am. J. Hum. Genet. 80, 393–406 (2007).

7. Domingo, A. et al. New insights into the genetics of X-linked dystonia-parkinsonism (XDP, DYT3). *Eur. J. Hum. Genet.* **23**, (2015).

8. Nolte, D., Niemann, S. & Müller, U. Specific sequence changes in multiple transcript system DYT3 are associated with X-linked dystonia parkinsonism. Proc. Natl. Acad. Sci. U. S. A. 100, 10347–10352 (2003).

9. Kitada, T. et al. Mutations in the parkin gene cause autosomal recessive juvenile parkinsonism. Nature 392, 605–608 (1998).

10. Kay, D. M. et al. A comprehensive analysis of deletions, multiplications, and copy number variations in PARK2. Neurology 75, 1189–1194 (2010).

11. Lesage, S. et al. Rare heterozygous parkin variants in French early-onset Parkinson disease patients and controls. J. Med. Genet. 45, (2008).

12. Daida, K. et al. Long-Read Sequencing Resolves a Complex Structural Variant in PRKN Parkinson’s Disease. Mov. Disord. 38, (2023).

13. Mahmoud, M. et al. Structural variant calling: the long and the short of it. Genome Biol 20, 246 (2019).

14. Kolmogorov, M. et al. Scalable Nanopore sequencing of human genomes provides a comprehensive view of haplotype-resolved variation and methylation. Nat. Methods 20, 1483–1492 (2023).

15. Smolka, M. et al. Detection of mosaic and population-level structural variants with Sniffles2. Nat. Biotechnol. 1–10 (2024).

16. Rizig, M. et al. Identification of genetic risk loci and causal insights associated with Parkinson’s disease in African and African admixed populations: a genome-wide association study. Lancet Neurol. 22, (2023).

17. Long, E. et al. The case for increasing diversity in tissue-based functional genomics datasets to understand human disease susceptibility. Nature communications 13, (2022).

18. Vitale, D. et al. GenoTools: An Open-Source Python Package for Efficient Genotype Data Quality Control and Analysis. bioRxiv 2024.03.26.586362 (2024) doi:10.1101/2024.03.26.586362.

19. Bray, S. M. et al. Signatures of founder effects, admixture, and selection in the Ashkenazi Jewish population. Proc. Natl. Acad. Sci. U. S. A. 107, 16222–16227 (2010).

20. Shafin, K. et al. Nanopore sequencing and the Shasta toolkit enable efficient de novo assembly of eleven human genomes. Nat. Biotechnol. 38, 1044–1053 (2020).

21. Krusche, P. et al. Best practices for benchmarking germline small variant calls in human genomes. (2018) doi:10.1101/270157.

22. Nalls, M. A. et al. Identification of novel risk loci, causal insights, and heritable risk for Parkinson’s disease: a meta-analysis of genome-wide association studies. Lancet Neurol. 18, 1091–1102 (2019).

23. Bellenguez, C. et al. New insights into the genetic etiology of Alzheimer’s disease and related dementias. Nat. Genet. 54, 412–436 (2022).

24. Chaisson, M. J. P. et al. Multi-platform discovery of haplotype-resolved structural variation in human genomes. Nat. Commun. 10, 1784 (2019).

25. Gustafson, J. A. et al. High-coverage nanopore sequencing of samples from the 1000 Genomes Project to build a comprehensive catalog of human genetic variation. Genome Res. (2024) doi:10.1101/gr.279273.124.

26. Beyter, D. et al. Long-read sequencing of 3,622 Icelanders provides insight into the role of structural variants in human diseases and other traits. Nat. Genet. 53, 779–786 (2021).

27. Mahmoud, M. et al. Utility of long-read sequencing for All of Us. Nat. Commun. 15, 837 (2024).

28. Collins, R. L. et al. A structural variation reference for medical and population genetics. Nature 581, 444–451 (2020).

29. Hickey, G. et al. Genotyping structural variants in pangenome graphs using the vg toolkit. Genome Biol. 21, 35 (2020).

30. Negi, S. et al. Advancing long-read nanopore genome assembly and accurate variant calling for rare disease detection. medRxiv (2024) doi:10.1101/2024.08.22.24312327.

31. Zheng, X., et al. STIX: Long-reads based Accurate Structural Variation Annotation at Population Scale. *bioRxiv* 2024.09.30.615931 (2024) doi:10.1101/2024.09.30.615931.

32. de Klein, N. et al. Brain expression quantitative trait locus and network analyses reveal downstream effects and putative drivers for brain-related diseases. Nat. Genet. 55, 377–388 (2023).

33. Raphael Gibbs, J., et al. Abundant Quantitative Trait Loci Exist for DNA Methylation and Gene Expression in Human Brain. PLoS Genet. 6, e1000952 (2010).

34. Dillman, A. A. et al. Transcriptomic profiling of the human brain reveals that altered synaptic gene expression is associated with chronological aging. Sci Rep 7, 16890 (2017).

35. Hoffman, G. E. et al. CommonMind Consortium provides transcriptomic and epigenomic data for Schizophrenia and Bipolar Disorder. Sci. Data 6, 180 (2019).

36. Chiang, C. et al. The impact of structural variation on human gene expression. Nat. Genet. 49, 692–699 (2017).

37. Genetic Architecture of Gene Expression in European and African Americans: An eQTL Mapping Study in GENOA. The American Journal of Human Genetics 106, 496–512 (2020).

38. Hormozdiari, F., Kostem, E., Kang, E. Y., Pasaniuc, B. & Eskin, E. Identifying causal variants at loci with multiple signals of association. Genetics 198, 497–508 (2014).

39. Kirsche, M. et al. Jasmine and Iris: Population-scale structural variant comparison and analysis. Nat. Methods 20, 408 (2023).

40. Mi, H. et al. Protocol Update for large-scale genome and gene function analysis with the PANTHER classification system (v.14.0). Nat. Protoc. 14, 703–721 (2019).

41. Rossi, M. N. et al. NLRP2 Regulates Proinflammatory and Antiapoptotic Responses in Proximal Tubular Epithelial Cells. Frontiers in Cell and Developmental Biology 7, 252 (2019).

42. Foox, J. et al. The SEQC2 epigenomics quality control (EpiQC) study. Genome Biol. 22, 332 (2021).

43. Jeong, H. et al. Evolution of DNA methylation in the human brain. Nat. Commun. 12, 2021 (2021).

44. Genner, R. et al. Assessing methylation detection for primary human tissue using Nanopore sequencing. bioRxiv 2024.02.29.581569 (2024) doi:10.1101/2024.02.29.581569.

45. *Methylation Clocks Do Not Predict Age or Alzheimer’s Disease Risk Across Genetically Admixed Individuals*.

46. The role of clustered protocadherins in neurodevelopment and neuropsychiatric diseases. Curr. Opin. Genet. Dev. 65, 144–150 (2020).

47. Galigniana, N. M. et al. Regulation of the glucocorticoid response to stress-related disorders by the Hsp90-binding immunophilin FKBP51. J. Neurochem. 122, 4–18 (2012).

48. Blair, L. J. et al. Accelerated neurodegeneration through chaperone-mediated oligomerization of tau. J. Clin. Invest. 123, 4158–4169 (2013).

49. Lunnon, K. et al. Methylomic profiling implicates cortical deregulation of ANK1 in Alzheimer’s disease. Nat. Neurosci. 17, 1164–1170 (2014).

50. Smith, R. G. et al. A meta-analysis of epigenome-wide association studies in Alzheimer’s disease highlights novel differentially methylated loci across cortex. Nat. Commun. 12, 3517 (2021).

51. Gardiner-Garden, M. & Frommer, M. CpG islands in vertebrate genomes. J. Mol. Biol. 196, 261–282 (1987).

52. Liu, W.-B. et al. LHX6 acts as a novel potential tumour suppressor with epigenetic inactivation in lung cancer. Cell Death Dis. 4, e882 (2013).

53. Yuan, F. et al. LHX6 is essential for the migration of human pluripotent stem cell-derived GABAergic interneurons. Protein Cell 11, 286–291 (2020).

54. Bahado-Singh, R. O. et al. Artificial intelligence and leukocyte epigenomics: Evaluation and prediction of late-onset Alzheimer’s disease. PLoS One 16, e0248375 (2021).

55. Humphries, C. et al. Alzheimer disease (AD) specific transcription, DNA methylation and splicing in twenty AD associated loci. Mol. Cell. Neurosci. 67, 37–45 (2015).

56. Aspra, Q. et al. Epigenome-wide analysis reveals DNA methylation alteration in ZFP57 and its target RASGFR2 in a Mexican population cohort with autism. Children (Basel) 9, 462 (2022).

57. Liu, Y., Toh, H., Sasaki, H., Zhang, X. & Cheng, X. An atomic model of Zfp57 recognition of CpG methylation within a specific DNA sequence. Genes Dev. 26, 2374–2379 (2012).

58. Razaghi, R. et al. Modbamtools: Analysis of single-molecule epigenetic data for long-range profiling, heterogeneity, and clustering. bioRxiv (2022) doi:10.1101/2022.07.07.499188.

59. Kuznetsov, N., et al. CNV-Finder: Streamlining Copy Number Variation Discovery. bioRxiv 2024.11.22.624040 (2024) doi:10.1101/2024.11.22.624040.

60. Liao, W.-W. et al. A Draft Human Pangenome Reference. (2022) doi:10.1101/2022.07.09.499321.

61. Stefansson, O. A. et al. The correlation between CpG methylation and gene expression is driven by sequence variants. Nat. Genet. 56, 1624–1631 (2024).

62. Kuschel, L. P. et al. Robust methylation-based classification of brain tumours using nanopore sequencing. Neuropathol. Appl. Neurobiol. 49, e12856 (2023).

63. Si, W. et al. Nanopore sequencing identifies differentially methylated genes in the central nervous system in experimental autoimmune encephalomyelitis. J. Neuroimmunol. 381, 578134 (2023).

64. Numata, S. et al. DNA methylation signatures in development and aging of the human prefrontal cortex. Am. J. Hum. Genet. 90, 260–272 (2012).

65. Kim, S. et al. DNA methylation associated with healthy aging of elderly twins. GeroScience 40, 469–484 (2018).

66. Gasparoni, G. et al. DNA methylation analysis on purified neurons and glia dissects age and Alzheimer’s disease-specific changes in the human cortex. Epigenetics Chromatin 11, (2018).

67. Sabbagh, J. J. et al. Age-associated epigenetic upregulation of the FKBP5 gene selectively impairs stress resiliency. PLoS One 9, e107241 (2014).

68. J Billingsley, K., et al. Processing human frontal cortex brain tissue for population-scale Oxford Nanopore long-read DNA sequencing SOP v2. (2022) doi:10.17504/protocols.io.kxygxzmmov8j/v2.

69. J Billingsley, K., et al. Processing human frontal cortex brain tissue for population-scale Oxford Nanopore long-read DNA sequencing SOP v2. (2022) doi:10.17504/protocols.io.kxygxzmmov8j/v2.

70. Baker, B. Processing human frontal cortex brain tissue for population-scale SQK-LSK114 Oxford Nanopore long-read DNA sequencing SOP v1. (2023) doi:10.17504/protocols.io.kxygx3zzog8j/v1.

71. Shafin, K. et al. Haplotype-aware variant calling with PEPPER-Margin-DeepVariant enables high accuracy in nanopore long-reads. Nat. Methods 18, 1322–1332 (2021).

72. Kolesnikov, A. et al. Local read haplotagging enables accurate long-read small variant calling. Nat. Commun. 15, 5907 (2024).

73. Poplin, R. et al. A universal SNP and small-indel variant caller using deep neural networks. Nat. Biotechnol. 36, 983–987 (2018).

74. Yun, T. et al. Accurate, scalable cohort variant calls using DeepVariant and GLnexus. Bioinformatics 36, 5582–5589 (2021).

75. Li, H. Minimap2: pairwise alignment for nucleotide sequences. Bioinformatics 34, 3094–3100 (2018).

76. Lorig-Roach, R. et al. Phased nanopore assembly with Shasta and modular graph phasing with GFAse. Genome Res. 34, 454–468 (2024).

77. English, A. C., Menon, V. K., Gibbs, R. A., Metcalf, G. A. & Sedlazeck, F. J. Truvari: refined structural variant comparison preserves allelic diversity. Genome Biol. 23, 1–20 (2022).

78. Danecek, P. et al. Twelve years of SAMtools and BCFtools. Gigascience 10, (2021).

79. Meylan, P., Dreos, R., Ambrosini, G., Groux, R. & Bucher, P. EPD in 2020: enhanced data visualization and extension to ncRNA promoters. Nucleic Acids Res. 48, D65–D69 (2020).

80. Website. https://www.ncbi.nlm.nih.gov/pmc/articles/PMC10659467/.

81. Patro, R., Duggal, G., Love, M. I., Irizarry, R. A. & Kingsford, C. Salmon provides fast and bias-aware quantification of transcript expression. Nat. Methods 14, 417–419 (2017).

82. Taylor-Weiner, A. et al. Scaling computational genomics to millions of individuals with GPUs. Genome Biol. 20, 228 (2019).

83. Vialle, R. A., de Paiva Lopes, K., Bennett, D. A., Crary, J. F. & Raj, T. Integrating whole-genome sequencing with multi-omic data reveals the impact of structural variants on gene regulation in the human brain. Nat. Neurosci. 25, (2022).

84. Billingsley, K. J. et al. Genome-Wide Analysis of Structural Variants in Parkinson Disease. Annals of neurology 93, (2023).

